# Gradient-based Optimization for mRNA Sequence Design

**DOI:** 10.1101/2025.10.22.683691

**Authors:** Hongmin Li, Goro Terai, Takumi Otagaki, Kiyoshi Asai

## Abstract

**Motivation:** Optimization of mRNA sequences presents fundamental challenges to balance multiple physicochemical/biological properties—including accessibility, stability, and translation efficiency—while preserving amino acid sequences. The discrete nature of RNA sequence design hinders direct application of gradient-based methods, despite their potential for leveraging modern deep learning predictors of the properties in biological sequence design.

**Results:** We present the Input Data Differentiable Designer (ID3) framework, a unified computational approach for mRNA sequence optimization that enables gradient-based optimization of discrete RNA sequences through innovative mathematical techniques. ID3 framework encompasses 12 constrained variants across four base configurations and three constraint mechanisms: Codon Profile Constraint, Amino Matching Softmax, and Lagrangian multipliers. The ID3 framework treats trained models as fixed differentiable functions while optimizing input data through continuous probability distributions. We also provide convergence analyses from the perspective of trained model input optimization.

**Availability and implementation:** https://github.com/Li-Hongmin/ID3.git

## 1. Introduction

Rational RNA sequence optimization underpins a wide range of transformative biomedical applications, from mRNA vaccines and protein replacement therapies to next-generation cancer immunotherapies (Pardi et al., 2018; Sample et al., 2019; Sahin et al., 2014). Yet, designing functional mRNA sequences remains a fundamental challenge in computational biology. Effective optimization must simultaneously balance multiple, often competing, biological objectives: ensuring efficient ribosome binding and translation initiation (Kozak, 1999), maintaining codon usage appropriate for the host organism (Sharp and Li, 1987), and preserving the encoded amino acid sequence required for correct protein function (Presnyak et al., 2015).

Codon optimization is particularly critical in practical applications, as synonymous codon choices can dramatically affect protein expression levels (Plotkin and Kudla, 2011), mRNA stability (Presnyak et al., 2015), and immunogenicity (Karikó et al., 2005)—factors that directly influence efficacy and safety of mRNA therapeutics (Pardi et al., 2018). However, traditional codon optimization methods suffer from fundamental limitations. Many rely on static lookup tables or hand-crafted heuristics, which are ill-suited to capture complex multi-objective trade-offs or adapt to diverse biological contexts (Reddy et al., 2015). Moreover, these approaches are difficult to integrate with modern deep learning models that predict mRNA accessibility or stability (Wayment-Steele et al., 2022). A central challenge arises from the inherently discrete nature of RNA sequences: conventional gradient-based optimization techniques require continuous and differentiable objective functions (Nocedal and Wright, 2006), creating a fundamental gap between powerful predictive models and practical sequence design.

This discrete-continuous optimization gap has limited the application of powerful neural network-based predictive models to RNA design. As a result, researchers have often been forced to rely on discrete optimization methods such as genetic algorithms and simulated annealing, which typically suffer from slow convergence and poor scalability (Zadeh et al., 2011; Hofacker et al., 1994).

Gradient-based approaches, such as activation maximization, have been developed to address vanishing gradient problems and have partially mitigated these challenges, improving convergence stability in molecular sequence design (Linder and Seelig, 2021). Furthermore, RNA folding models based on a differentiable partition function have shown that gradient-based optimization with probabilistic sequence representations can simultaneously optimize thermodynamic stability and codon adaptation index (CAI) (Krueger and Ward, 2025). However, these methods still lack theoretical convergence guarantees and do not incorporate systematic codon-constraint mechanisms to ensure amino acid sequence preservation during optimization. Finally, deep generative approaches for codon optimization (Li et al., 2024) require training new models with codon-level token inputs. This additional training requirement limits the direct reuse of high-accuracy mRNA property prediction models that were originally trained on nucleotide sequences.

To address these limitations—including the discrete-continuous optimization gap, the lack of codon-level constraints in gradient-based optimization, and the difficulty of reusing existing predictive models—we introduce the Input Data Differentiable Designer (ID3) framework. ID3 employs a decoupled optimization-evaluation architecture that separates the optimization process from candidate evaluation (Fig. 1). The optimization track transforms sequence logits into probabilities and supports two modes: *Soft mode* uses continuous probabilities for gradient optimization, while *Hard mode* uses discretized sequences with Straight-Through Estimator (STE) gradients. The evaluation track independently collects sequence-prediction pairs at every iteration, enabling models to accept probability distributions as input while maintaining biological validity through discrete candidate evaluation.

**Fig. 1:**
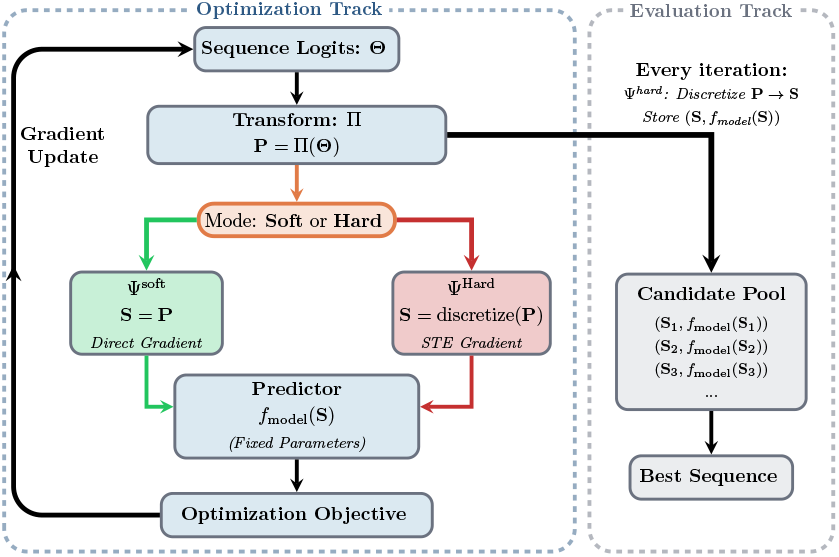
ID3 Framework Overview. The decoupled optimization-evaluation architecture showing the optimization track (Soft/Hard modes) and the independent evaluation track for biological validity assessment.

**Fig. 2:**
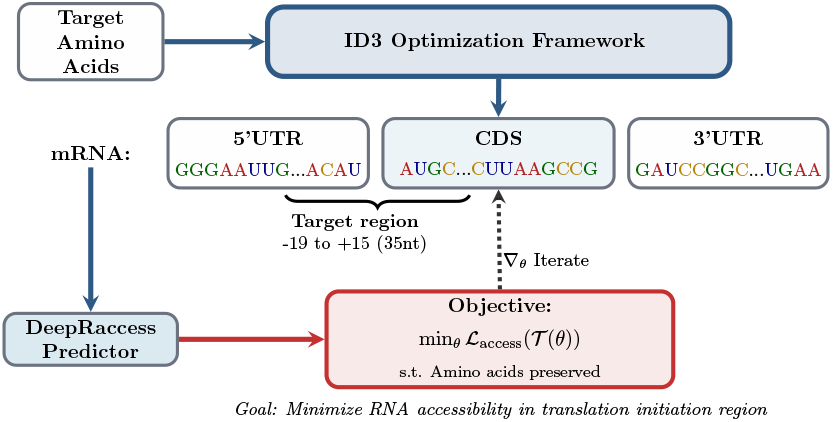
DeepRaccess Integration Framework. The ID3 optimization framework integrates with DeepRaccess predictor to minimize RNA accessibility in the translation initiation region while maintaining target amino acid sequences. The workflow shows the optimization pipeline from target amino acids through mRNA design to accessibility prediction and iterative refinement via gradient updates.

By treating pre-trained predictors as fixed differentiable functions and optimizing over continuous probability distributions, ID3 integrates with existing deep learning models without requiring retraining. This formulation provides theoretical convergence guarantees and enables performance improvements in RNA design tasks such as mRNA accessibility and joint CAI optimization.

## 2. Methods

### 2.1. The mathematical framework of ID3

The ID3 provides a systematic approach to mRNA sequence optimization. A discrete RNA sequence is represented by a matrix of learnable continuous parameters Θ. The Θ is transformed to a matrix of probabilities **P** by Π, and the **P** is converted to a ‘generalized sequence’ **S** by Ψ. The objective function *L* is computed by a fixed pre-trained model *f*_model_(*T* (Θ)) (*T* = Π ○ Ψ). The core mathematical objective of the optimization is:

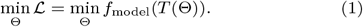

The Θ is optimized for ℒ by the gradient-based updates ∇:

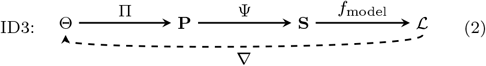

Fig. 1 shows the decoupled optimization-evaluation architecture where the optimization track supports both Soft mode (continuous gradient optimization) and Hard mode (discrete sequence optimization via STE), while the evaluation track independently collects and evaluates candidates every iteration.

#### 2.1.1. Transformation Functions and ID3 Variants

The ID3 framework defines two key transformation functions that progressively convert learnable parameters to sequences through probability distributions, where Θ represents the matrix of learnable parameters for a sequence.

##### Parameter-to-Probability Transformation (Π)

This transformation converts the parameter matrix Θ to a probability matrix **P** representing the probabilities at each position of the sequence, using Gumbel-Softmax (Jang et al., 2016; Gumbel, 1958):

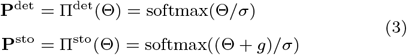

where *g* is a sampled noise from Gumbel distribution, and *σ* is the temperature parameter.

##### Probability-to-Sequence Transformation (Ψ)

This transformation converts probability distributions **P** to generalized sequences **S**, supporting both soft and hard modes:

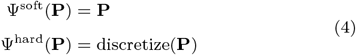

where discretize(**P**) converts probability distributions to valid discrete sequences through constraint-aware strategies.

These transformations enable two mutually exclusive optimization modes: the Soft mode, which performs stable gradient optimization over continuous probability distributions, and the Hard mode, which applies the Straight-Through Estimator (STE) (Bengio et al., 2013) to discrete sequences to maintain gradient flow.

The unified transformation function *T* is defined as:

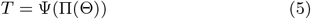

where Π ∈ { Π^det^, Π^sto^ } and Ψ ∈ {Ψ^soft^, Ψ^hard^ }, yielding four base operational modes as detailed in Table 1:

**Table 1.** ID3 Base Operational Modes.

| Mode | Parameter Type | Output Type | Notation |
| --- | --- | --- | --- |
| Det.Soft | Deterministic | Soft | $T^{\text{det.soft}}(\Theta)$ |
| Det.Hard | Deterministic | Hard | $T^{\text{det.hard}}(\Theta)$ |
| Sto.Soft | Stochastic | Soft | $T^{\text{sto.soft}}(\Theta)$ |
| Sto.Hard | Stochastic | Hard | $T^{\text{sto.hard}}(\Theta)$ |

The deterministic/stochastic distinction is expressed through the choice of probability transformation Π^det/sto^, while the soft/hard distinction occurs through the sequence transformation Ψ^soft/hard^. This separation allows constraint mechanisms to focus on their core functionality without duplicating mode-specific behavior.

These four base operational modes from Table 1 combine with three constraint mechanisms to create 12 constrained variants for comprehensive biological sequence optimization.

### 2.2. Biological Constraints and Convergence Analysis

#### 2.2.1. Codon Constraint Challenge in mRNA Design

The central challenge in computational mRNA optimization lies in balancing two competing requirements: optimizing RNA sequence properties while maintaining the exact amino acid sequence needed for protein function. This arises from codon degeneracy—most amino acids can be encoded by multiple synonymous codons, each affecting different RNA properties yet preserving protein function.

##### Mathematical Problem Formulation

Given a target amino acid sequence **y** = [*y*_1_, *y*_2_, …, *y*_*M*_] where *M* is the number of amino acids, we seek to optimize RNA properties while maintaining perfect amino acid sequence preservation:

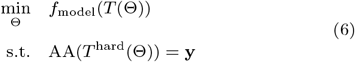

Here Θ ∈ ℝ^*L×*4^ represents learnable RNA parameters with *L* = 3*M*, *f*_model_ is the fixed trained predictor, and AA(·) denotes the amino acid sequence function that requires discrete input. Constraints are enforced through three mechanisms: Codon Profile Constraint, Amino Matching Softmax, and Lagrangian multipliers.

#### 2.2.2. Three Constraint Integration Methods

The ID3 framework supports amino acid constraint preservation through three distinct integration approaches:

##### (1) Codon Profile Constraint

This approach fundamentally redesigns the parameter space to operate directly on codon probabilities rather than nucleotide probabilities. The method achieves constraint satisfaction by construction—ensuring amino acid constraint satisfaction since invalid codons are excluded from the parameter space.

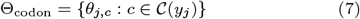

where each parameter corresponds only to codons that encode the target amino acid. The probability computation ensures only valid codons receive non-zero probabilities:

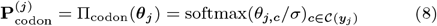

This approach reduces parameter dimensionality from *L ×* 4 to 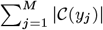, typically yielding 40-60% parameter reduction while providing the strongest constraint guarantee.

##### (2) Amino Matching Softmax

This method preserves the full RNA parameter space while ensuring amino acid constraints through similarity-based projection. Codon probabilities are determined by computing inner product between RNA parameters and codon encodings as the slimilarity.

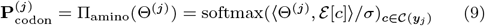

where Θ^(*j*)^ is the RNA parameter vector for the codon of *j*-th amino acid, and ℰ [*c*] is the one-hot encoding of codon *c*. The inner product ⟨ Θ^(*j*)^, *ℰ* [*c*] ⟩ measures similarity between RNA parameters and valid codons, effectively projecting RNA parameters onto the codon space. This approach ensures that codons more similar to the RNA parameters receive higher probabilities, while restricting to valid codons only. The softmax normalization guarantees that probabilities sum to 1 across valid codons only. Unlike Codon Profile, this approach allows gradient flow through the full RNA space while projecting only at the probability level, providing a balance between expressiveness and constraint satisfaction.

##### (3) Lagrangian Multiplier

This approach adopts the classical optimization strategy of soft penalty constraints, operating on the full RNA parameter space without structural modifications. The constraint violation is measured using L2 distance to the closest valid codon encoding:

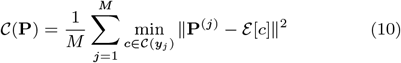

The total optimization objective combines the original model loss with the constraint penalty:

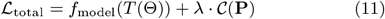

where *λ* is the Lagrangian multiplier that controls the penalty strength. The optimization objective becomes minimizing ℒ_total_, balancing model performance with constraint adherence. The Lagrangian multiplier is adaptively updated using subgradient method with decaying step size to ensure convergence:

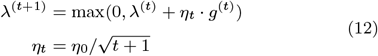

where *g*^(*t*)^ is the subgradient of the constraint at iteration *t*, and *η*_*t*_ is the decaying step size. This soft constraint approach allows temporary constraint violations during optimization while gradually enforcing compliance, providing the most flexible optimization landscape but requiring careful tuning of the penalty coefficient.

#### 2.2.3. Soft and Hard Mode Transformation Patterns

After constraint integration, we employ transformation functions to convert codon probabilities into continuous probability distributions (soft mode) or discrete sequences (hard mode). Both Codon Profile and Amino Matching constraints operate on codon probability space **P**_codon_, applying *R*_nuc_(**P**_codon_) for soft mode and Onehot(**P**_codon_) for hard mode, where:

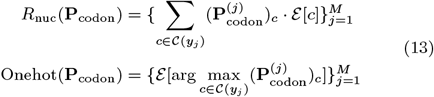

Their fundamental difference lies in probability generation: Codon Profile directly parameterizes codon space, while Amino Matching projects RNA parameters through similarity matching. In contrast, Lagrangian operates on RNA probability space **P**, using **P** directly in soft mode but requiring Π_amino_ projection before discretization in hard mode. Table 2 summarizes these patterns.

**Table 2.** Soft and Hard Mode Transformation Patterns.

| Constraint Type | $\Psi^{\text{soft}}$ | $\Psi^{\text{hard}}$ |
| --- | --- | --- |
| Codon Profile | $R_{\text{nuc}}(\mathbf{P}_{\text{codon}})$ | $\text{Onehot}(\mathbf{P}_{\text{codon}})$ |
| Amino Matching | $R_{\text{nuc}}(\mathbf{P}_{\text{codon}})$ | $\text{Onehot}(\mathbf{P}_{\text{codon}})$ |
| Lagrangian | $\mathbf{P}$ | $\text{Onehot}(\Pi_{\text{amino}}(\mathbf{P}))$ |

#### 2.2.4. Theoretical Convergence Guarantees

We provide, to our knowledge, the first theoretical convergence analysis for sequence optimization from the trained model input optimization perspective. We mathematically prove that the ID3 framework achieves convergence guarantees based on certain assumptions about predictor model quality. These assumptions can typically be considered as satisfied in ID3 practical applications. The theoretical framework systematically covers all 12 ID3 variants through hierarchical theorems with detailed proofs in Supplementary Section S5, providing rigorous theoretical foundation for practical RNA sequence design applications.

### 2.3. Applications

To demonstrate the ID3 framework’s optimization performance and multi-objective extension capability, we conducted comprehensive evaluations on two complementary tasks: mRNA accessibility optimization and mRNA accessibility-CAI joint optimization.

#### 2.3.1. Accessibility Optimization

RNA accessibility represents the thermodynamic energy required to make a region accessible, measured in kcal/mol. Lower accessibility values indicate more easily accessed regions, facilitating ribosome binding and translation initiation (Bernhart et al., 2011). We use DeepRaccess (Hara et al., 2023), a deep learning model that accelerates the classical Raccess algorithm, as the accessibility predictor *f*_access_ in the ID3 framework.

##### Model Architecture Integration

Without requiring architectural modifications or fine-tuning of trained models, the ID3 framework extends token-based neural architectures like DeepRaccess to support probabilistic sequences through a soft embedding mechanism that enables gradient-based optimization:

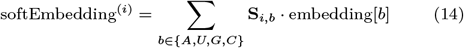

where **S**_*i,b*_ *∈* [0, 1] represents the probability of nucleotide *b* at position *i* from the ID3 sequence representation, and embedding[*b*] ∈ ℝ^*d*^ is the trained embedding vector for nucleotide *b*.

##### mRNA Structure and CDS Focus

The framework targets CDS optimization within the complete mRNA structure:

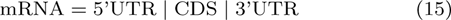

##### Loss Function Design

The ribosome binding site and start codon region from position −19 to +15 relative to the ATG start codon determines translation initiation efficiency (Terai and Asai, 2020). We reformulate the general constrained optimization framework (Eq. (6)) for the accessibility optimization task as:

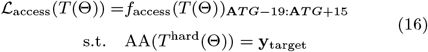

where the objective function focuses on minimizing RNA accessibility in the critical translation initiation region, and the amino acid constraint is enforced through one of the three constraint mechanisms described in Section 2.2.

#### 2.3.2. Accessibility-CAI Joint Optimization

The Codon Adaptation Index (CAI) (Sharp and Li, 1987) measures codon usage bias relative to highly expressed genes in the target organism, reflecting the degree of adaptation to organism-specific translational machinery. To enable simultaneous optimization of RNA accessibility and CAI, we extend Eq. (16) through two complementary technical modifications:

##### CAI-aware Discretization

Our core approach enhances the discretization process to consider both accessibility and CAI objectives for both soft and hard modes. The optimal sequence is found by solving:

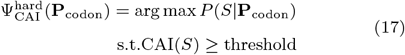

where *S* is the discrete sequence. The CAI threshold is set based on organism-specific requirements (e.g., 0.8 for *E. coli*). To solve this constrained optimization, we introduce a linear interpolation of the codon probability distribution with the target organism’s codon usage frequency vector:

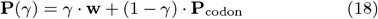

where **w** is the codon usage frequency vector for the target organism, and *γ* ∈ [0, 1] controls the trade-off between model probability and codon adaptation. By adjusting *γ*, we can find a sequence that maximizes *P* (*S* **P**(*γ*)) while satisfying the CAI constraint. We solve this constrained optimization problem through a binary search-based method that iteratively finds the optimal sequence by balancing probability consistency and CAI requirements. The detailed algorithm is in Supplementary Section S7. For the Lagrangian constraint mechanism, this approach requires first converting RNA probabilities to codon probabilities using Π_amino_ before applying the CAI enhancement.

##### Multi-objective Joint Loss Option

As an additional enhancement option for both soft and hard modes, CAI constraints can also be integrated through multi-objective joint loss:

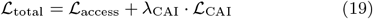

where ℒ_CAI_ measures deviation from the target CAI threshold. For probabilistic optimization with codon distributions, we compute the expected CAI as a weighted average of codon weights according to their probabilities, enabling gradient-based optimization of the CAI penalty term (see Supplementary Section S8.1 for formulation). The penalty coefficient *λ*_CAI_ controls the relative importance of the two objectives, offering flexible multi-objective optimization.

## 3. Results

### 3.1. Computational Experiment Setup

We evaluated ID3 across 12 diverse proteins (64-1,273 amino acids) with 20 independent runs per protein-variant combination. The dataset spans multiple functional categories and organisms (Table 3) for comprehensive framework assessment. The UTR sequences are derived from the pET-11a expression vector (Supplementary Section S9).

**Table 3.** Protein dataset.

| UniProt ID | Protein Name | Organism | Length (aa) |
| --- | --- | --- | --- |
| O15263 | Defensin beta 4A | *H. sapiens* | 64 |
| P99999 | Cytochrome c | *H. sapiens* | 105 |
| P00004 | Cytochrome c | *E. caballus* | 105 |
| P01308 | Insulin | *H. sapiens* | 110 |
| P01825 | Immunoglobulin heavy chain | *H. sapiens* | 116 |
| P31417 | Fatty acid-binding protein 2 | *M. sexta* | 132 |
| P61626 | Lysozyme C | *H. sapiens* | 148 |
| P42212 | Green fluorescent protein | *A. victoria* | 238 |
| P04637 | Tumor suppressor p53 | *H. sapiens* | 393 |
| P0DTC9 | SARS-CoV-2 Nucleoprotein | *SARS-CoV-2* | 419 |
| P0CG48 | Polyubiquitin-C | *H. sapiens* | 685 |
| P0DTC2 | SARS-CoV-2 Spike protein | *SARS-CoV-2* | 1273 |

We conducted comprehensive computational experiments across two optimization scenarios: accessibility-only optimization (144 experiments across 12 variants × 12 proteins) focusing solely on minimizing mRNA accessibility, and CAI joint optimization incorporating additional codon adaptation constraints. For

Accessibility-CAI joint optimization, we set the CAI target threshold to 0.8. For fair comparison, all methods used exactly 1000 DeepRaccess evaluations per run. For performance validation, we implemented baseline optimization algorithms: Genetic Algorithm (GA) with population size 20 and 50 generations (20×50=1000 evaluations), and Simulated Annealing (SA) with exponential cooling schedule over 1000 iterations. We also evaluated partial optimization variants (GA-10, SA-10) that optimize only the first 10 codons in the translation initiation region while assigning remaining codons to the most frequent codon for each amino acid, and obtained the theoretical 10-codon optimum (C10) through combinatorial exhaustive search. Our ID3 framework optimizes one sequence per run, while baseline methods explore multiple candidates per iteration. All methods were evaluated across the same 12 proteins with 20 independent runs each, totaling 480 baseline experiments.

### 3.2. Performance Analysis

Our computational evaluation establishes Amino.Sto.Soft as the optimal configuration, achieving 0.929 kcal/mol accessibility scores. Remarkably, 10 of 12 ID3 variants outperform all baseline methods including the combinatorial 10-codon optimum (C10: 1.354 kcal/mol), with the best variant achieving 1.6× improvement over C10, 2.3× over GA (2.118 kcal/mol), and 3.7× over SA (2.776 kcal/mol) (Fig. 3).

**Fig. 3:**
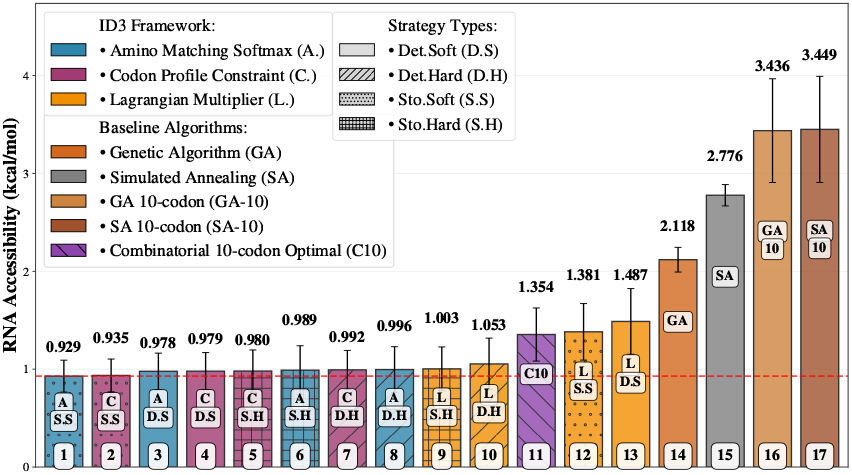
Performance Ranking of ID3 Variants and Baseline Methods. ID3 framework dominates with 11 variants (ranks 1-10) outperforming all baselines. Combinatorial 10-codon Optimal (C10) ranks 11th, followed by Lagrangian.Sto.Soft (rank 12), traditional full-sequence methods (GA, SA: ranks 14-15), and partial optimization approaches (GA-10, SA-10: ranks 16-17). Amino.Sto.Soft achieves optimal performance at 0.926 kcal/mol.

CAI joint optimization computational experiments reveal accessibility-adaptation trade-offs, with hard constraint variants achieving superior performance in multi-objective scenarios (Fig. 4). Amino.Sto.Hard with CAI penalty achieves optimal performance (0.977 kcal/mol), while soft constraint methods show degraded performance under codon adaptation requirements. Performance analysis shows no significant difference between Codon Profile and Amino Matching constraint mechanisms (see Supplementary Section S2.3), while stochastic hard strategies consistently outperform soft approaches in CAI joint optimization (Figures 5 and 6).

**Fig. 4:**
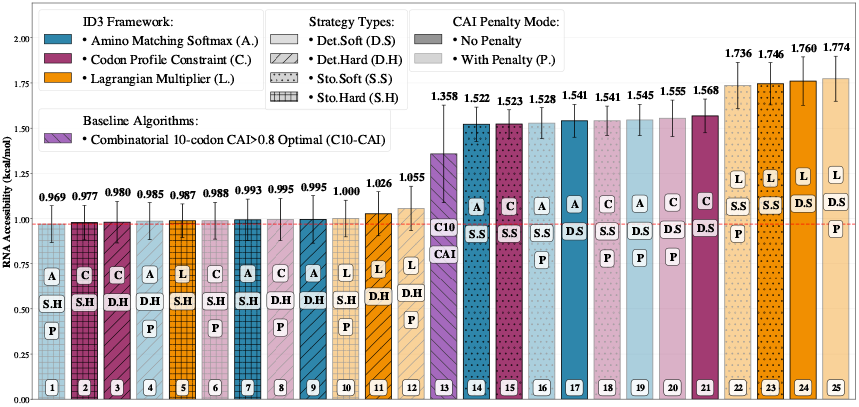
Performance Ranking of ID3 Variants for CAI Joint Optimization. Hard constraint variants achieve superior performance in multi-objective scenarios, with Amino.Sto.Hard+Penalty optimal at 0.969 kcal/mol. The combinatorial 10-codon CAI*>*0.8 optimum (C10-CAI) ranks 13th at 1.358 kcal/mol.

**Fig. 5:**
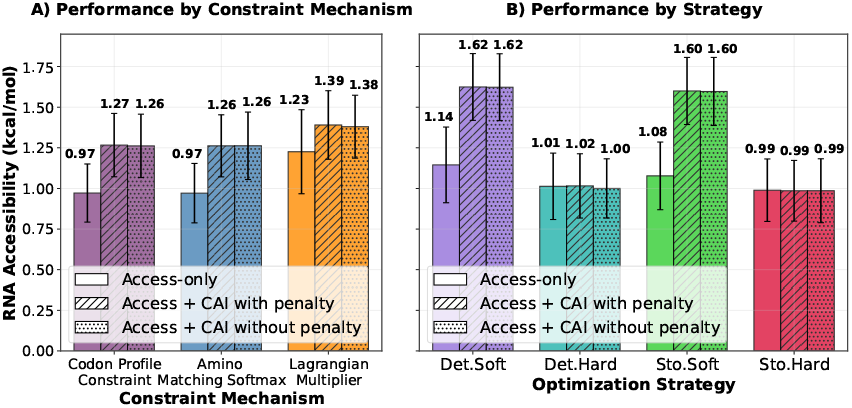
Performance by Constraint and Strategy. Codon Profile and Amino Matching achieve similar performance. Stochastic strategies (Sto.Soft, Sto.Hard) outperform deterministic approaches.

**Fig. 6:**
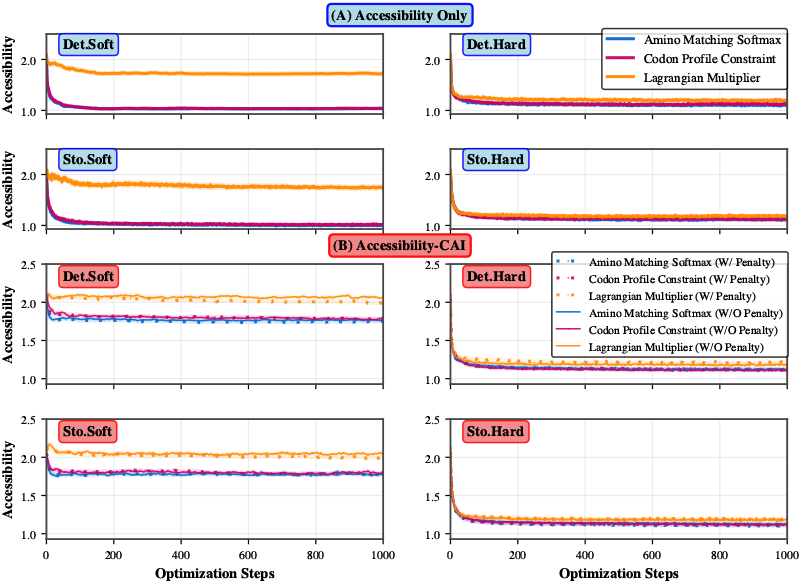
Convergence Analysis by Strategy Type. Comparison across constraint mechanisms and strategy variants The upper panel shows accessibility-only optimization with three convergence trajectories and comparable performance between soft and hard constraints. The lower panel reveals CAI joint optimization where hard constraint variants consistently outperform soft variants.

Fig. 6 reveals a fundamental insight into single-objective versus multi-objective optimization dynamics. The accessibility-only optimization (upper panel) demonstrates that soft and hard constraint variants achieve comparable performance, indicating effective exploration within the unified optimization landscape where accessibility serves as the singular objective. However, CAI joint optimization (lower panel) alters this dynamic: the introduction of penalty-based objectives creates target misalignment between accessibility improvement and codon adaptation, causing soft constraint methods to become trapped in suboptimal regions due to conflicting gradient signals. In contrast, hard constraint variants using Straight-Through Estimation maintain optimization efficacy by preserving exact constraint satisfaction without penalty-induced gradient interference, explaining their superior performance in multi-objective scenarios. This is confirmed by computational experiments showing Amino.Sto.Hard achieving optimal CAI joint optimization performance (0.977 kcal/mol) compared to Amino.Sto.Soft variants ranking significantly lower. This observation establishes that constraint satisfaction mechanisms become important when optimization objectives exhibit spatial or temporal misalignment in the parameter space. Notably, ID3’s gradient-based optimization surpasses even exhaustive combinatorial search: the 10-codon optimum (C10: 1.459 kcal/mol) required evaluating all 1,364,736 possible combinations, yet 11 of 12 ID3 variants achieve better performance through intelligent full-sequence optimization. The performance gap between full-sequence (GA: 2.118) and partial optimization (GA-10: 3.436) further confirms the importance of global sequence context in RNA design.

Detailed numerical results and statistical comparisons are in Supplementary Section S1 and S2.

### 3.3. Case Study: O15263 Protein Optimization

We demonstrate framework effectiveness through O15263 (Defensin beta 4A) optimization using Amino.Sto.Soft. The case study reveals dynamic progression of nucleotide probabilities, rapid convergence of sequence probabilities and loss values to stable states, and systematic AU content balancing throughout the optimization process (Fig. 7). The observed convergence behavior validates the framework’s ability to identify global optima efficiently, supporting practical applicability for therapeutic mRNA design.

**Fig. 7:**
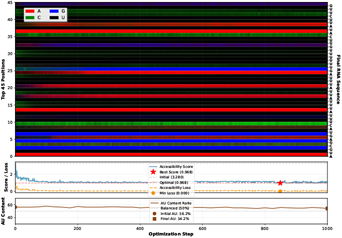
Case Study: O15263 Optimization Trajectory. Dynamic optimization trajectory showing nucleotide probability progression (top), rapid convergence of accessibility from 3.28 to 0.97 kcal/mol (middle), and AU content balancing throughout the process (bottom).

## 4. Conclusion

We present the ID3 framework, which bridges the discrete–continuous optimization gap through a decoupled optimization–evaluation architecture. ID3 separates continuous probability optimization from discrete sequence evaluation, allowing models to accept probabilistic inputs while preserving amino acid sequences. This design removes the need for model retraining or architectural modification, enabling direct use of existing predictors with probabilistic inputs.

The framework systematically supports 12 constrained variants formed by combining four operational modes (deterministic/stochastic and soft/hard) with three distinct mathematical constraint mechanisms for amino acid preservation, and establishes theoretical convergence guarantees from the trained model input optimization perspective. Through computational evaluations on mRNA accessibility optimization and Accessibility-CAI joint optimization across diverse protein sequences, ID3 demonstrates superior performance over traditional optimization methods, establishing a principled and practical foundation for gradient-based optimization of discrete biological sequences.

## References

Y. Bengio, N. Léonard, and A. Courville. Estimating or propagating gradients through stochastic neurons for conditional computation. arXiv preprint arXiv:1308.3432, 2013.

S. H. Bernhart, I. L. Hofacker, and P. F. Stadler. Rna accessibility in cubic time. Algorithms for Molecular Biology, 6:3, 2011. doi: 10.1186/1748-7188-6-3.

E. J. Gumbel. Statistics of extremes. Columbia University Press., 1958.

K. Hara, N. Iwano, T. Fukunaga, and M. Hamada. Deepraccess: high-speed rna accessibility prediction using deep learning. Frontiers in Bioinformatics, 3:1275787, 2023. doi: 10.3389/fbinf.2023.1275787.

L. Hofacker, W. Fontana, P. F. Stadler, L. S. Bonhoeffer, M. Tacker, and P. Schuster. Fast folding and comparison of rna secondary structures. Monatshefte für Chemie/Chemical Monthly, 125(2): 167–188, 1994.

E. Jang, S. Gu, and B. Poole. Categorical reparameterization with gumbel-softmax. arXiv preprint arXiv:1611.01144, 2016.

K. Karikó, M. Buckstein, H. Ni, and D. Weissman. Suppression of rna recognition by toll-like receptors: the impact of nucleoside modification and the evolutionary origin of rna. Immunity, 23 (2):165–175, 2005.

M. Kozak. Initiation of translation in prokaryotes and eukaryotes. Gene, 234(2): 187–208, 1999.

R. K. Krueger and M. Ward. JAX-RNAfold: scalable differentiable folding. Bioinformatics, 41(5):btaf203, 04 2025. doi: 10.1093/bioinformatics/btaf203.

Y. Li, F. Wang, J. Yang, Z. Han, L. Chen, W. Jiang, H. Zhou, T. Li, Z. Tang, J. Deng, X. He, G. Zha, J. Hu, Y. Hu, L. Wu, C. Zhan, C. Sun, Y. He, and Z. Xie. Deep generative optimization of mrna codon sequences for enhanced protein production and therapeutic efficacy. bioRxiv, 2024. doi: 10.1101/2024.09.06.611590.

J. Linder and G. Seelig. Fast activation maximization for molecular sequence design. BMC Bioinformatics, 22(1): 510, 2021. ISSN 1471-2105. doi: 10.1186/s12859-021-04437-5.

J. Nocedal and S. Wright. Numerical optimization. Springer Science & Business Media., 2006.

N. Pardi, M. J. Hogan, F. W. Porter, and D. Weissman. mrna vaccines—a new era in vaccinology. Nature Reviews Drug Discovery, 17(4): 261–279, 2018.

J. B. Plotkin and G. Kudla. Synonymous but not the same: the causes and consequences of codon bias. Nature Reviews Genetics, 12(1): 32–42, 2011.

V. Presnyak, N. Alhusaini, Y. H. Chen, S. Martin, N. Morris, N. Kline, S. Olson, D. Weinberg, K. E. Baker, B. R. Graveley, and J. Coller. Codon optimality is a major determinant of mrna stability. Cell, 160(6): 1111–1124, 2015.

P. S. Reddy, S. Mahanty, T. Kaul, S. Nair, S. K. Sopory, and M. K. Reddy. Plant codon usage and the prediction of gene expression. Molecular Genetics and Genomics, 290(4): 1335–1348, 2015.

U. Sahin, K. Karikó, and O. Tureci. mrna-based therapeutics—developing a new class of drugs. Nature Reviews Drug Discovery, 13(10): 759–780, 2014.

P. J. Sample, B. Wang, D. W. Reid, V. Presnyak, I. J. McFadyen, D. R. Morris, and G. Seelig. Human 5’ utr design and variant effect prediction from a massively parallel translation assay. Nature Biotechnology, 37(7): 803–809, 2019.

P. M. Sharp and W. H. Li. The codon adaptation index-a measure of directional synonymous codon usage bias, and its potential applications. Nucleic Acids Research, 15(3): 1281–1295, 1987.

G. Terai and K. Asai. Improving the prediction accuracy of protein abundance in escherichia coli using mrna accessibility. Nucleic Acids Research, 48(14):e81, 06 2020. doi: 10.1093/nar/gkaa481.

H. K. Wayment-Steele, W. Wu, I. I. Kallemeijn, M. Khorshid, K. T. Beier, and R. Das. Deep learning models for predicting rna degradation via dual crowdsourcing. bioRxiv, 2022., 2022.

J. N. Zadeh, C. D. Steenberg, J. S. Bois, B. R. Wolfe, M. B. Pierce, A. R. Khan, R. M. Dirks, and N. A. Pierce. Nupack: Analysis and design of nucleic acid systems. Journal of Computational Chemistry, 32(1): 170–173, 2011.

