## Supplementary material for "Gradient-based Optimization for mRNA Sequence Design": ID3_Supplementary_Material

*Bioinformatics*, 2025, 1–25
doi: 10.1093/bioinformatics/example-supp
Advance Access Publication Date: Day Month Year
Computational Biology

### Supplementary Material: Gradient-based Optimization for mRNA Sequence Design

**Li Hongmin,[1,*] Goro Terai,[1] Takumi Otagaki[1] and Kiyoshi Asai[1]**

[1]Department of Computational Biology and Medical Sciences, Graduate School of Frontier Sciences, University of Tokyo, 5-1-5 Kashiwanoha, 277-8561, Kashiwa-shi, Chiba, Japan

[*]Corresponding author.[†]To whom correspondence should be addressed.; ORCID: 0000-0002-1059-2519[‡]Correspondence may also be addressed to Kiyoshi Asai.

FOR PUBLISHER ONLY Received on Date Month Year; revised on Date Month Year; accepted on Date Month Year

#### Abstract

**This supplementary material provides comprehensive theoretical analysis, methodological details, and computational validation for the ID3 framework for mRNA sequence design.**

**Key words:** convergence analysis, optimization theory, mRNA design, CAI optimization, constraint mechanisms, statistical validation

#### S1. Detailed Performance Results

This section provides comprehensive numerical results for all computational evaluations referenced in the main paper.

##### S1.1. Complete Performance Comparison Table

Table S1 presents the detailed numerical results for all 12 ID3 variants compared with baseline optimization algorithms (GA and SA) across 12 diverse proteins. Each cell shows mean accessibility scores with standard deviations from 20 independent runs.

**Table 1.** Performance comparison of ID3 variants and baseline algorithms for RNA accessibility optimization (kcal/mol, $mean_{std}$)

| Method | O15263 | P00004 | P01308 | P01825 | P04637 | P0CG48 | P0DTC2 | P0DTC9 | P31417 | P42212 | P61626 | P99999 | Avg | Rank |
|---|---|---|---|---|---|---|---|---|---|---|---|---|---|---|
| ***Baseline Methods*** | | | | | | | | | | | | | | |
| Genetic Algorithm | $1.900_{0.14}$ | $1.278_{0.09}$ | $2.440_{0.18}$ | $1.901_{0.10}$ | $2.454_{0.14}$ | $2.314_{0.08}$ | $2.680_{0.10}$ | $2.360_{0.12}$ | $2.172_{0.12}$ | $2.034_{0.11}$ | $2.620_{0.14}$ | $1.266_{0.10}$ | 2.118 | 14 |
| Simulated Annealing | $2.570_{0.17}$ | $1.910_{0.10}$ | $3.289_{0.11}$ | $2.636_{0.16}$ | $3.036_{0.07}$ | $2.807_{0.07}$ | $3.113_{0.04}$ | $2.915_{0.09}$ | $3.040_{0.08}$ | $2.717_{0.09}$ | $3.358_{0.15}$ | $1.924_{0.10}$ | 2.776 | 15 |
| Genetic Algorithm (10-codon) | $3.563_{0.004}$ | $2.789_{0.002}$ | $4.480_{0.005}$ | $3.184_{0.005}$ | $3.481_{0.001}$ | $3.124_{0.001}$ | $3.582_{0.000}$ | $3.548_{0.000}$ | $3.970_{0.003}$ | $3.203_{0.003}$ | $4.565_{0.005}$ | $2.887_{0.003}$ | 3.531 | 16 |
| Simulated Annealing (10-codon) | $3.583_{0.009}$ | $2.791_{0.003}$ | $4.515_{0.01}$ | $3.194_{0.006}$ | $3.486_{0.003}$ | $3.125_{0.001}$ | $3.583_{0.000}$ | $3.549_{0.001}$ | $3.982_{0.005}$ | $3.210_{0.005}$ | $4.587_{0.009}$ | $2.895_{0.005}$ | 3.542 | 17 |
| Combinatorial 10-codon Optimal | 1.256 | 0.987 | 1.822 | 1.521 | 1.657 | 1.079 | 1.645 | 1.538 | 1.093 | 1.270 | 1.218 | 1.159 | 1.354 | 11 |
| ***ID3 Framework Variants*** | | | | | | | | | | | | | | |
| Codon.Sto.Soft | $1.030_{0.09}$ | **$0.778_{0.01}$** | $1.227_{0.02}$ | $1.167_{0.04}$ | $1.012_{0.04}$ | $0.748_{0.01}$ | $0.912_{0.02}$ | $0.751_{0.05}$ | $1.060_{0.03}$ | $0.882_{0.03}$ | $0.917_{0.07}$ | $0.731_{0.003}$ | 0.935 | 2 |
| Codon.Sto.Hard | $1.299_{0.39}$ | $0.905_{0.06}$ | $1.222_{0.08}$ | $1.193_{0.07}$ | $0.992_{0.04}$ | $0.806_{0.07}$ | $1.038_{0.17}$ | $0.814_{0.09}$ | $0.966_{0.07}$ | **$0.815_{0.05}$** | $0.833_{0.07}$ | $0.858_{0.10}$ | 0.978 | 4 |
| Codon.Det.Soft | $1.117_{0.09}$ | $0.793_{0.02}$ | $1.254_{0.02}$ | $1.295_{0.05}$ | $1.043_{0.02}$ | $0.778_{0.02}$ | $0.933_{0.02}$ | $0.766_{0.05}$ | $1.138_{0.05}$ | $0.909_{0.04}$ | $1.000_{0.04}$ | $0.736_{0.004}$ | 0.980 | 5 |
| Codon.Det.Hard | $1.247_{0.30}$ | $0.924_{0.06}$ | $1.247_{0.09}$ | $1.235_{0.09}$ | $1.029_{0.06}$ | $0.809_{0.04}$ | $1.113_{0.15}$ | $0.788_{0.06}$ | $0.990_{0.09}$ | $0.823_{0.08}$ | $0.859_{0.09}$ | $0.853_{0.08}$ | 0.993 | 8 |
| Amino.Sto.Soft | **$0.998_{0.02}$** | $0.778_{0.01}$ | $1.245_{0.02}$ | **$1.156_{0.05}$** | **$0.989_{0.05}$** | **$0.747_{0.009}$** | **$0.891_{0.02}$** | **$0.740_{0.02}$** | $1.018_{0.06}$ | $0.910_{0.03}$ | $0.945_{0.06}$ | **$0.730_{0.004}$** | 0.929 | 1 |
| Amino.Det.Soft | $1.053_{0.06}$ | $0.795_{0.02}$ | $1.270_{0.01}$ | $1.281_{0.04}$ | $1.033_{0.02}$ | $0.762_{0.008}$ | $0.940_{0.02}$ | $0.757_{0.06}$ | $1.117_{0.08}$ | $0.941_{0.03}$ | $1.043_{0.05}$ | $0.735_{0.01}$ | 0.977 | 3 |
| Amino.Sto.Hard | $1.439_{0.48}$ | $0.922_{0.05}$ | $1.214_{0.15}$ | $1.181_{0.10}$ | $1.008_{0.07}$ | $0.808_{0.03}$ | $1.011_{0.14}$ | $0.791_{0.05}$ | **$0.960_{0.09}$** | $0.817_{0.04}$ | **$0.831_{0.10}$** | $0.861_{0.08}$ | 0.987 | 6 |
| Amino.Det.Hard | $1.302_{0.42}$ | $0.914_{0.07}$ | **$1.188_{0.15}$** | $1.223_{0.08}$ | $1.009_{0.09}$ | $0.834_{0.04}$ | $1.045_{0.15}$ | $0.781_{0.06}$ | $1.043_{0.18}$ | $0.819_{0.06}$ | $0.846_{0.07}$ | $0.885_{0.08}$ | 0.991 | 7 |
| Lagrangian.Sto.Hard | $1.356_{0.26}$ | $0.802_{0.01}$ | $1.240_{0.14}$ | $1.280_{0.10}$ | $1.041_{0.06}$ | $0.805_{0.04}$ | $1.047_{0.13}$ | $0.859_{0.08}$ | $1.061_{0.10}$ | $0.880_{0.08}$ | $0.852_{0.09}$ | $0.799_{0.04}$ | 1.002 | 9 |
| Lagrangian.Det.Hard | $1.650_{0.39}$ | $0.868_{0.09}$ | $1.319_{0.08}$ | $1.261_{0.11}$ | $1.042_{0.07}$ | $0.793_{0.05}$ | $1.002_{0.11}$ | $0.930_{0.08}$ | $1.167_{0.15}$ | $0.881_{0.05}$ | $0.887_{0.10}$ | $0.866_{0.11}$ | 1.055 | 10 |
| Lagrangian.Sto.Soft | $1.513_{0.15}$ | $1.010_{0.08}$ | $1.964_{0.13}$ | $1.635_{0.05}$ | $1.380_{0.15}$ | $1.031_{0.06}$ | $1.472_{0.20}$ | $1.258_{0.10}$ | $1.474_{0.21}$ | $1.342_{0.12}$ | $1.264_{0.11}$ | $1.078_{0.11}$ | 1.368 | 12 |
| Lagrangian.Det.Soft | $1.680_{0.35}$ | $1.059_{0.11}$ | $2.091_{0.18}$ | $1.692_{0.08}$ | $1.516_{0.16}$ | $1.086_{0.06}$ | $1.672_{0.23}$ | $1.343_{0.10}$ | $1.583_{0.20}$ | $1.481_{0.15}$ | $1.380_{0.14}$ | $1.144_{0.11}$ | 1.477 | 13 |

*Note: Bold values indicate lowest (best) score per protein. Rankings based on average performance.*

##### S1.2. Accessibility-CAI Results

This section provides detailed results for Accessibility-CAI computational experiments, including both CAI scores achieved and corresponding accessibility performance. Each variant is evaluated under two CAI integration methods: penalty-based (multi-objective joint loss) and no-penalty (CAI-aware discretization).

© The Author 2025. Published by Oxford University Press. All rights reserved. For permissions, please

##### S1.2.1. Combined Accessibility Performance

Table S2 shows the accessibility performance for all 12 variants under both CAI integration methods. Each variant displays two rows: the default row shows performance without penalty (CAI-aware discretization method), and the +Penalty row shows performance with penalty-based integration (multi-objective joint loss approach). Rankings are calculated across all 24 entries (12 variants × 2 modes).

**Table 2.** Accessibility-CAI joint optimization performance (accessibility, kcal/mol, $mean_{std}$)

| Method | O15263 | P00004 | P01308 | P01825 | P04637 | P0CG48 | P0DTC2 | P0DTC9 | P31417 | P42212 | P61626 | P99999 | Avg | Rank |
|---|---|---|---|---|---|---|---|---|---|---|---|---|---|---|
| *Baseline Methods* | | | | | | | | | | | | | | |
| Combinatorial 10-codon Optimal | 1.302 | 0.987 | 1.822 | 1.521 | 1.657 | 1.079 | 1.645 | 1.538 | 1.093 | 1.270 | 1.218 | 1.159 | 1.358 | 13 |
| *ID3 Framework Variants* | | | | | | | | | | | | | | |
| Codon.Det.Soft | $1.811_{0.11}$ | $1.247_{0.11}$ | $1.948_{0.09}$ | $1.507_{0.06}$ | $1.639_{0.09}$ | $1.270_{0.08}$ | $1.704_{0.15}$ | $1.646_{0.07}$ | $1.575_{0.07}$ | $1.425_{0.06}$ | $1.589_{0.11}$ | $1.452_{0.11}$ | 1.568 | 21 |
| Codon.Det.Soft +Penalty | $1.778_{0.10}$ | $1.257_{0.11}$ | $1.909_{0.07}$ | $1.504_{0.04}$ | $1.567_{0.11}$ | $1.267_{0.11}$ | $1.698_{0.16}$ | $1.618_{0.11}$ | $1.601_{0.07}$ | $1.364_{0.08}$ | $1.629_{0.11}$ | $1.464_{0.13}$ | 1.555 | 20 |
| Codon.Det.Hard | $1.230_{0.31}$ | $0.930_{0.06}$ | $1.195_{0.23}$ | $1.224_{0.08}$ | $1.004_{0.07}$ | $0.803_{0.04}$ | $0.995_{0.13}$ | $0.811_{0.07}$ | $1.032_{0.16}$ | $0.836_{0.06}$ | $0.837_{0.09}$ | $0.859_{0.09}$ | 0.980 | 3 |
| Codon.Det.Hard +Penalty | $1.339_{0.44}$ | $0.937_{0.05}$ | $1.255_{0.26}$ | $1.230_{0.06}$ | $1.015_{0.04}$ | $0.804_{0.05}$ | $0.987_{0.14}$ | $0.798_{0.05}$ | $0.997_{0.11}$ | $0.818_{0.05}$ | $0.862_{0.07}$ | $0.893_{0.08}$ | 0.995 | 8 |
| Codon.Sto.Soft | $1.752_{0.08}$ | $1.186_{0.11}$ | $1.874_{0.07}$ | $1.508_{0.04}$ | $1.578_{0.06}$ | $1.143_{0.08}$ | $1.777_{0.03}$ | $1.645_{0.07}$ | $1.569_{0.08}$ | $1.310_{0.07}$ | $1.566_{0.10}$ | $1.366_{0.14}$ | 1.523 | 15 |
| Codon.Sto.Soft +Penalty | $1.783_{0.12}$ | $1.265_{0.06}$ | $1.908_{0.08}$ | $1.514_{0.03}$ | $1.560_{0.07}$ | $1.197_{0.10}$ | $1.767_{0.08}$ | $1.619_{0.08}$ | $1.594_{0.06}$ | $1.315_{0.09}$ | $1.583_{0.07}$ | $1.386_{0.12}$ | 1.541 | 18 |
| Codon.Sto.Hard | $1.293_{0.32}$ | $0.903_{0.06}$ | $1.226_{0.11}$ | $1.185_{0.10}$ | $1.010_{0.05}$ | $0.802_{0.05}$ | $\mathbf{0.976_{0.12}}$ | $\mathbf{0.779_{0.06}}$ | $1.002_{0.09}$ | $0.853_{0.06}$ | $0.820_{0.07}$ | $0.879_{0.07}$ | 0.977 | 2 |
| Codon.Sto.Hard +Penalty | $1.320_{0.32}$ | $0.898_{0.06}$ | $1.184_{0.13}$ | $1.239_{0.09}$ | $\mathbf{0.987_{0.06}}$ | $0.808_{0.05}$ | $1.033_{0.15}$ | $0.787_{0.06}$ | $1.038_{0.11}$ | $0.825_{0.05}$ | $0.861_{0.08}$ | $0.872_{0.07}$ | 0.988 | 6 |
| Amino.Det.Soft | $1.816_{0.09}$ | $1.242_{0.12}$ | $1.886_{0.07}$ | $1.546_{0.05}$ | $1.550_{0.09}$ | $1.278_{0.06}$ | $1.771_{0.08}$ | $1.626_{0.13}$ | $1.522_{0.11}$ | $1.349_{0.08}$ | $1.613_{0.13}$ | $1.289_{0.09}$ | 1.541 | 17 |
| Amino.Det.Soft +Penalty | $1.800_{0.10}$ | $1.235_{0.11}$ | $1.869_{0.07}$ | $1.499_{0.05}$ | $1.579_{0.09}$ | $1.286_{0.04}$ | $1.781_{0.11}$ | $1.629_{0.08}$ | $1.556_{0.10}$ | $1.328_{0.07}$ | $1.608_{0.09}$ | $1.375_{0.11}$ | 1.545 | 19 |
| Amino.Det.Hard | $1.387_{0.49}$ | $0.892_{0.07}$ | $\mathbf{1.180_{0.14}}$ | $\mathbf{1.172_{0.13}}$ | $1.014_{0.08}$ | $0.814_{0.04}$ | $1.084_{0.15}$ | $0.812_{0.07}$ | $0.990_{0.14}$ | $0.853_{0.08}$ | $0.869_{0.11}$ | $0.872_{0.08}$ | 0.995 | 9 |
| Amino.Det.Hard +Penalty | $1.214_{0.27}$ | $0.923_{0.05}$ | $1.201_{0.13}$ | $1.227_{0.16}$ | $1.041_{0.07}$ | $0.829_{0.03}$ | $1.041_{0.14}$ | $0.803_{0.06}$ | $0.985_{0.08}$ | $0.835_{0.08}$ | $0.848_{0.09}$ | $0.875_{0.07}$ | 0.985 | 4 |
| Amino.Sto.Soft | $1.746_{0.12}$ | $1.204_{0.15}$ | $1.836_{0.12}$ | $1.530_{0.06}$ | $1.557_{0.10}$ | $1.255_{0.05}$ | $1.768_{0.09}$ | $1.596_{0.11}$ | $1.536_{0.13}$ | $1.309_{0.06}$ | $1.643_{0.06}$ | $1.281_{0.10}$ | 1.522 | 14 |
| Amino.Sto.Soft +Penalty | $1.772_{0.08}$ | $1.244_{0.10}$ | $1.867_{0.09}$ | $1.521_{0.04}$ | $1.541_{0.07}$ | $1.250_{0.07}$ | $1.749_{0.08}$ | $1.646_{0.11}$ | $1.523_{0.12}$ | $1.311_{0.04}$ | $1.573_{0.11}$ | $1.337_{0.11}$ | 1.528 | 16 |
| Amino.Sto.Hard | $1.487_{0.49}$ | $0.898_{0.07}$ | $1.237_{0.11}$ | $1.214_{0.14}$ | $0.997_{0.05}$ | $0.805_{0.04}$ | $0.995_{0.09}$ | $0.787_{0.07}$ | $\mathbf{0.975_{0.09}}$ | $\mathbf{0.796_{0.04}}$ | $0.874_{0.10}$ | $0.847_{0.08}$ | 0.993 | 7 |
| Amino.Sto.Hard +Penalty | $\mathbf{1.187_{0.32}}$ | $0.911_{0.07}$ | $1.208_{0.10}$ | $1.187_{0.14}$ | $0.997_{0.06}$ | $\mathbf{0.802_{0.06}}$ | $1.015_{0.12}$ | $0.799_{0.06}$ | $0.987_{0.08}$ | $0.831_{0.05}$ | $0.831_{0.08}$ | $0.872_{0.07}$ | 0.969 | 1 |
| Lagrangian.Det.Soft | $1.828_{0.11}$ | $1.454_{0.12}$ | $2.044_{0.15}$ | $1.689_{0.05}$ | $1.766_{0.10}$ | $1.459_{0.09}$ | $1.953_{0.19}$ | $1.767_{0.14}$ | $1.916_{0.19}$ | $1.969_{0.21}$ | $1.691_{0.10}$ | $1.590_{0.15}$ | 1.760 | 24 |
| Lagrangian.Det.Soft +Penalty | $1.853_{0.10}$ | $1.431_{0.13}$ | $2.014_{0.09}$ | $1.675_{0.09}$ | $1.710_{0.12}$ | $1.447_{0.10}$ | $2.084_{0.04}$ | $1.834_{0.12}$ | $1.997_{0.27}$ | $1.890_{0.23}$ | $1.721_{0.09}$ | $1.625_{0.12}$ | 1.774 | 25 |
| Lagrangian.Det.Hard | $1.333_{0.30}$ | $0.833_{0.08}$ | $1.270_{0.19}$ | $1.282_{0.10}$ | $1.036_{0.08}$ | $0.810_{0.05}$ | $1.135_{0.14}$ | $0.920_{0.10}$ | $1.169_{0.21}$ | $0.872_{0.07}$ | $0.846_{0.06}$ | $0.809_{0.08}$ | 1.026 | 11 |
| Lagrangian.Det.Hard +Penalty | $1.453_{0.28}$ | $0.851_{0.09}$ | $1.389_{0.18}$ | $1.288_{0.12}$ | $1.022_{0.06}$ | $0.805_{0.06}$ | $1.149_{0.22}$ | $0.941_{0.08}$ | $1.198_{0.18}$ | $0.894_{0.06}$ | $0.836_{0.07}$ | $0.840_{0.08}$ | 1.055 | 12 |
| Lagrangian.Sto.Soft | $1.837_{0.15}$ | $1.484_{0.11}$ | $1.985_{0.06}$ | $1.677_{0.06}$ | $1.679_{0.13}$ | $1.491_{0.07}$ | $1.967_{0.11}$ | $1.810_{0.11}$ | $1.789_{0.16}$ | $1.966_{0.24}$ | $1.718_{0.07}$ | $1.543_{0.13}$ | 1.746 | 23 |
| Lagrangian.Sto.Soft +Penalty | $1.839_{0.13}$ | $1.430_{0.12}$ | $2.032_{0.13}$ | $1.646_{0.08}$ | $1.746_{0.12}$ | $1.507_{0.06}$ | $2.022_{0.10}$ | $1.796_{0.12}$ | $1.798_{0.23}$ | $1.774_{0.22}$ | $1.720_{0.08}$ | $1.528_{0.15}$ | 1.736 | 22 |
| Lagrangian.Sto.Hard | $1.279_{0.25}$ | $\mathbf{0.783_{0.02}}$ | $1.230_{0.13}$ | $1.285_{0.10}$ | $1.023_{0.08}$ | $0.806_{0.05}$ | $1.063_{0.14}$ | $0.859_{0.09}$ | $1.077_{0.10}$ | $0.865_{0.06}$ | $\mathbf{0.812_{0.07}}$ | $\mathbf{0.768_{0.03}}$ | 0.987 | 5 |
| Lagrangian.Sto.Hard +Penalty | $1.384_{0.27}$ | $0.800_{0.02}$ | $1.213_{0.16}$ | $1.294_{0.12}$ | $1.006_{0.07}$ | $0.804_{0.05}$ | $1.095_{0.18}$ | $0.834_{0.08}$ | $1.076_{0.11}$ | $0.883_{0.06}$ | $0.827_{0.08}$ | $0.781_{0.02}$ | 1.000 | 10 |

*Note: Each ID3 variant shows two rows: default (No Penalty) and +Penalty. C10 = Combinatorial 10-codon optimal with CAI¿0.8 constraint. Bold = best per protein. First 20 runs.*

##### S1.2.2. Combined CAI Scores

Table S3 presents the CAI scores achieved by each variant under both integration methods, demonstrating the effectiveness of both penalty-based and discretization-based CAI integration approaches. The table follows the same format as Table S2, with each variant showing two rows for the two integration methods.

**Table 3.** Accessibility-CAI joint optimization scores (CAI values, $mean_{std}$)

| Method | O15263 | P00004 | P01308 | P01825 | P04637 | P0CG48 | P0DTC2 | P0DTC9 | P31417 | P42212 | P61626 | P99999 | Avg |
|---|---|---|---|---|---|---|---|---|---|---|---|---|---|
| *Baseline Methods* | | | | | | | | | | | | | |
| Combinatorial 10-codon Optimal | 0.803 | 0.903 | 0.878 | 0.892 | 0.979 | 0.979 | 0.990 | 0.978 | 0.920 | 0.945 | 0.891 | 0.924 | 0.923 |
| *ID3 Framework Variants* | | | | | | | | | | | | | |
| Codon.Det.Soft | $0.809_{0.008}$ | $0.807_{0.01}$ | $0.804_{0.003}$ | $0.807_{0.010}$ | $0.801_{0.001}$ | $0.800_{0.000}$ | $0.800_{0.000}$ | $0.805_{0.006}$ | $0.804_{0.003}$ | $0.803_{0.002}$ | $0.802_{0.003}$ | $0.806_{0.005}$ | 0.804 |
| Codon.Det.Soft +Penalty | $0.809_{0.009}$ | $0.806_{0.005}$ | $0.804_{0.003}$ | $0.802_{0.002}$ | $0.801_{0.002}$ | $0.800_{0.000}$ | $0.800_{0.000}$ | $0.802_{0.002}$ | $0.804_{0.004}$ | $0.804_{0.006}$ | $0.806_{0.007}$ | $0.806_{0.008}$ | 0.804 |
| Codon.Det.Hard | $0.807_{0.007}$ | $0.804_{0.002}$ | $0.803_{0.003}$ | $0.804_{0.002}$ | $0.801_{0.002}$ | $0.800_{0.000}$ | $0.801_{0.002}$ | $0.803_{0.003}$ | $0.802_{0.002}$ | $0.803_{0.003}$ | $0.805_{0.004}$ | $0.803_{0.004}$ | 0.803 |
| Codon.Det.Hard +Penalty | $0.809_{0.009}$ | $0.805_{0.003}$ | $0.803_{0.003}$ | $0.805_{0.004}$ | $0.803_{0.004}$ | $0.800_{0.000}$ | $0.800_{0.000}$ | $0.804_{0.008}$ | $0.802_{0.002}$ | $0.804_{0.005}$ | $0.805_{0.004}$ | $0.804_{0.003}$ | 0.804 |
| Codon.Sto.Soft | $0.808_{0.009}$ | $0.805_{0.005}$ | $0.804_{0.003}$ | $0.802_{0.002}$ | $0.801_{0.001}$ | $0.800_{0.000}$ | $0.800_{0.000}$ | $0.803_{0.006}$ | $0.803_{0.002}$ | $0.805_{0.006}$ | $0.803_{0.003}$ | $0.805_{0.007}$ | 0.803 |
| Codon.Sto.Soft +Penalty | $0.807_{0.008}$ | $0.804_{0.004}$ | $0.806_{0.004}$ | $0.802_{0.003}$ | $0.802_{0.003}$ | $0.800_{0.000}$ | $0.800_{0.000}$ | $0.801_{0.002}$ | $0.803_{0.002}$ | $0.802_{0.003}$ | $0.804_{0.005}$ | $0.804_{0.004}$ | 0.803 |
| Codon.Sto.Hard | $0.806_{0.006}$ | $0.805_{0.004}$ | $0.802_{0.002}$ | $0.802_{0.002}$ | $0.801_{0.001}$ | $0.800_{0.000}$ | $0.800_{0.001}$ | $0.803_{0.006}$ | $0.802_{0.002}$ | $0.801_{0.002}$ | $0.803_{0.002}$ | $0.803_{0.003}$ | 0.802 |
| Codon.Sto.Hard +Penalty | $0.807_{0.007}$ | $0.803_{0.003}$ | $0.805_{0.009}$ | $0.803_{0.003}$ | $0.801_{0.001}$ | $0.800_{0.001}$ | $0.800_{0.000}$ | $0.801_{0.001}$ | $0.801_{0.001}$ | $0.801_{0.001}$ | $0.803_{0.003}$ | $0.803_{0.003}$ | 0.802 |
| Amino.Det.Soft | $0.812_{0.01}$ | $0.809_{0.009}$ | $0.805_{0.004}$ | $0.809_{0.01}$ | $0.801_{0.001}$ | $0.801_{0.001}$ | $0.800_{0.000}$ | $0.808_{0.01}$ | $0.806_{0.004}$ | $0.803_{0.003}$ | $0.805_{0.005}$ | $0.807_{0.006}$ | 0.806 |
| Amino.Det.Soft +Penalty | $0.806_{0.006}$ | $0.806_{0.006}$ | $0.807_{0.006}$ | $0.807_{0.009}$ | $0.802_{0.002}$ | $0.801_{0.001}$ | $0.801_{0.004}$ | $0.802_{0.002}$ | $0.805_{0.004}$ | $0.805_{0.006}$ | $0.804_{0.004}$ | $0.809_{0.009}$ | 0.805 |
| Amino.Det.Hard | $0.809_{0.006}$ | $0.805_{0.004}$ | $0.805_{0.003}$ | $0.804_{0.003}$ | $0.802_{0.003}$ | $0.801_{0.001}$ | $0.800_{0.000}$ | $0.804_{0.004}$ | $0.803_{0.003}$ | $0.805_{0.008}$ | $0.805_{0.003}$ | $0.804_{0.004}$ | 0.804 |
| Amino.Det.Hard +Penalty | $0.809_{0.006}$ | $0.807_{0.005}$ | $0.804_{0.002}$ | $0.806_{0.005}$ | $0.803_{0.002}$ | $0.801_{0.001}$ | $0.801_{0.000}$ | $0.802_{0.001}$ | $0.803_{0.002}$ | $0.805_{0.008}$ | $0.804_{0.002}$ | $0.804_{0.003}$ | 0.804 |
| Amino.Sto.Soft | $0.809_{0.008}$ | $0.807_{0.006}$ | $0.806_{0.006}$ | $0.805_{0.004}$ | $0.802_{0.001}$ | $0.801_{0.001}$ | $0.801_{0.001}$ | $0.801_{0.002}$ | $0.804_{0.003}$ | $0.804_{0.006}$ | $0.805_{0.004}$ | $0.807_{0.006}$ | 0.804 |
| Amino.Sto.Soft +Penalty | $0.807_{0.009}$ | $0.808_{0.008}$ | $0.805_{0.007}$ | $0.804_{0.004}$ | $0.802_{0.002}$ | $0.801_{0.001}$ | $0.800_{0.000}$ | $0.802_{0.004}$ | $0.804_{0.005}$ | $0.802_{0.002}$ | $0.804_{0.003}$ | $0.806_{0.006}$ | 0.804 |
| Amino.Sto.Hard | $0.807_{0.006}$ | $0.804_{0.004}$ | $0.805_{0.003}$ | $0.805_{0.006}$ | $0.802_{0.002}$ | $0.801_{0.001}$ | $0.801_{0.000}$ | $0.804_{0.003}$ | $0.803_{0.003}$ | $0.804_{0.002}$ | $0.804_{0.004}$ | $0.804_{0.004}$ | 0.804 |
| Amino.Sto.Hard +Penalty | $0.806_{0.005}$ | $0.805_{0.004}$ | $0.803_{0.003}$ | $0.806_{0.004}$ | $0.802_{0.001}$ | $0.800_{0.000}$ | $0.801_{0.000}$ | $0.804_{0.004}$ | $0.803_{0.003}$ | $0.804_{0.003}$ | $0.804_{0.003}$ | $0.803_{0.003}$ | 0.803 |
| Lagrangian.Det.Soft | $0.812_{0.01}$ | $0.810_{0.010}$ | $0.808_{0.007}$ | $0.804_{0.005}$ | $0.803_{0.003}$ | $0.801_{0.001}$ | $0.802_{0.002}$ | $0.806_{0.01}$ | $0.803_{0.003}$ | $0.805_{0.006}$ | $0.826_{0.03}$ | $0.808_{0.009}$ | 0.807 |
| Lagrangian.Det.Soft +Penalty | $0.810_{0.01}$ | $0.806_{0.005}$ | $0.807_{0.008}$ | $0.807_{0.007}$ | $0.804_{0.004}$ | $0.802_{0.002}$ | $0.803_{0.003}$ | $0.802_{0.002}$ | $0.806_{0.005}$ | $0.804_{0.003}$ | $0.806_{0.007}$ | $0.807_{0.004}$ | 0.805 |
| Lagrangian.Det.Hard | $0.809_{0.008}$ | $0.804_{0.004}$ | $0.807_{0.008}$ | $0.803_{0.002}$ | $0.801_{0.000}$ | $0.803_{0.008}$ | $0.800_{0.000}$ | $0.801_{0.003}$ | $0.803_{0.003}$ | $0.801_{0.000}$ | $0.803_{0.004}$ | $0.805_{0.005}$ | 0.803 |
| Lagrangian.Det.Hard +Penalty | $0.809_{0.007}$ | $0.805_{0.004}$ | $0.806_{0.006}$ | $0.804_{0.004}$ | $0.802_{0.002}$ | $0.802_{0.005}$ | $0.800_{0.000}$ | $0.801_{0.002}$ | $0.802_{0.002}$ | $0.802_{0.002}$ | $0.803_{0.004}$ | $0.803_{0.003}$ | 0.803 |
| Lagrangian.Sto.Soft | $0.811_{0.01}$ | $0.809_{0.009}$ | $0.808_{0.006}$ | $0.806_{0.008}$ | $0.802_{0.003}$ | $0.801_{0.001}$ | $0.802_{0.002}$ | $0.803_{0.003}$ | $0.802_{0.003}$ | $0.804_{0.004}$ | $0.814_{0.02}$ | $0.810_{0.009}$ | 0.806 |
| Lagrangian.Sto.Soft +Penalty | $0.808_{0.009}$ | $0.805_{0.006}$ | $0.806_{0.008}$ | $0.807_{0.007}$ | $0.802_{0.002}$ | $0.801_{0.001}$ | $0.801_{0.000}$ | $0.802_{0.002}$ | $0.803_{0.002}$ | $0.802_{0.002}$ | $0.803_{0.002}$ | $0.806_{0.005}$ | 0.804 |
| Lagrangian.Sto.Hard | $0.806_{0.006}$ | $0.804_{0.003}$ | $0.805_{0.004}$ | $0.803_{0.003}$ | $0.801_{0.001}$ | $0.800_{0.000}$ | $0.800_{0.000}$ | $0.801_{0.000}$ | $0.803_{0.003}$ | $0.801_{0.001}$ | $0.802_{0.003}$ | $0.804_{0.003}$ | 0.803 |
| Lagrangian.Sto.Hard +Penalty | $0.807_{0.006}$ | $0.804_{0.003}$ | $0.804_{0.003}$ | $0.803_{0.002}$ | $0.801_{0.000}$ | $0.800_{0.000}$ | $0.800_{0.000}$ | $0.801_{0.001}$ | $0.801_{0.001}$ | $0.801_{0.001}$ | $0.802_{0.002}$ | $0.803_{0.003}$ | 0.802 |

*Note: Each ID3 variant shows two rows: default (No Penalty) and +Penalty. C10 = Combinatorial 10-codon optimal (CAI scores at CAI¿0.8 constrained sequences). First 20 runs.*

#### S2. Computational Experiment Setup and Results Analysis

##### S2.1. Supplementary Computational Experiment Setup

This section details the configurations and implementations for all baseline algorithms evaluated in this study. All computational experiments use normalized datasets with exactly 20 independent runs per configuration to ensure statistical reliability.

**S2.1.1. Experimental Scope**

The experimental design encompasses 9,600 optimization runs across ID3 framework variants and baseline algorithms, plus 1,364,736 C10 combinatorial evaluations. Table 4 summarizes the complete experimental scope, including three categories of baseline methods: full-sequence optimization (GA, SA), partial-sequence optimization (GA-10, SA-10), and exhaustive combinatorial search (C10).

**Table 4.** Experimental Design and Scope

| Category | Configuration | Runs |
|---|---|---|
| **ID3 Framework** | | |
| Accessibility | 12 proteins × 3 constraints × 4 variants × 20 runs | 2,880 |
| CAI (Penalty) | 12 proteins × 3 constraints × 4 variants × 20 runs | 2,880 |
| CAI (No-Penalty) | 12 proteins × 3 constraints × 4 variants × 20 runs | 2,880 |
| *ID3 Subtotal* | | *8,640* |
| **Baseline Algorithm** | | |
| Full-sequence (GA, SA) | 12 proteins × 2 methods × 20 runs | 480 |
| 10-codon (GA-10, SA-10) | 12 proteins × 2 methods × 20 runs | 480 |
| C10 Combinatorial | Exhaustive search: 1,364,736 combinations (6,144–331,776 per protein) | Theoretical |
| *Baseline Subtotal* | | *960 + C10* |
| **Total Runs** | | **9,600** |

**S2.1.2. Baseline Algorithm Implementation**

All baseline algorithms preserve the target amino acid sequence through synonymous codon substitutions only. The objective function minimizes DeepRaccess accessibility scores for the ATG−19:+15 region across 12 proteins.

Full-Sequence Optimization (GA, SA)

These methods optimize all codon positions throughout the entire sequence. Both algorithms were configured to use exactly 1000 DeepRaccess evaluations per run for fair comparison.

**Genetic Algorithm.** The GA employs a population of 20 individuals evolving over 49 generations (yielding $20 \times 50 = 1000$ total evaluations). Tournament selection (size 3) identifies parents for reproduction. Offspring are generated through uniform crossover at the codon level (rate 0.8) and synonymous codon mutation (rate 0.1, affecting approximately 10% of optimizable codons). Elitism preserves the top 2 individuals across generations.

**Simulated Annealing.** The SA performs 999 iterations plus 1 initial evaluation. Temperature decreases exponentially from $T_{\text{init}} = 5.0$ to $T_{\text{final}} = 0.01$ with cooling rate $\alpha = (T_{\text{final}}/T_{\text{init}})^{1/999} \approx 0.9938$. Each iteration randomly replaces 2 codons with synonymous alternatives. Metropolis acceptance criterion $P(\text{accept}) = \exp(-\Delta E/T)$ governs uphill moves. To escape local minima, the algorithm restarts with elevated temperature ($2 \times T_{\text{init}}$) after 100 iterations without improvement (maximum 3 restarts).

Partial-Sequence Optimization (GA-10, SA-10)

These constrained variants optimize only the first 10 codons following the start codon (positions 1-10), while fixing all remaining codons to the highest-probability synonymous codon from E.coli BL21(DE3) codon usage table. This design tests whether localized optimization can achieve comparable performance to full-sequence methods.

The GA-10 and SA-10 variants use identical algorithmic parameters as their full-sequence counterparts (population size 20, 49 generations for GA-10; 999 iterations for SA-10), but restrict all operations (crossover, mutation, neighbor generation) to the 10-codon optimization window. Each method was evaluated with 20 independent runs per protein (240 total runs per method).

Combinatorial Exhaustive Search (C10)

The C10 baseline represents the theoretical optimum achievable through exhaustive combinatorial search over a limited codon window. This method enumerates all possible synonymous codon combinations for the first 10 codons following the start codon, fixing remaining positions to default codons.

The search space comprises 1,364,736 total combinations across 12 proteins, ranging from 6,144 to 331,776 combinations per protein depending on synonymous degeneracy. For each protein, all combinations are evaluated to identify the global optimum within this constrained 10-codon subspace. This baseline establishes an upper bound on partial-sequence optimization performance and enables direct comparison between gradient-based full-sequence methods (ID3) and exhaustive local search.

#### S2.2. Results Analysis

**S2.2.1. Variant Performance Rankings**

We evaluated 12 ID3 variants in accessibility and 24 variants in Accessibility-CAI. Complete rankings are provided in Tables S1 and S2. The following analysis focuses on statistically validated performance tiers and identifies mode-specific patterns.

Accessibility

Table 5 summarizes pairwise statistical tests comparing top-ranked variants. Complete performance rankings for all 12 variants appear in Table S1.

**Table 5.** Pairwise Statistical Tests for Top Variant Rankings (Accessibility)

| Comparison | Variants (Rank, Mean kcal/mol) | Δ | p-value | Sig? |
|---|---|---|---|---|
| Rank 1 vs 2 | Amino.Sto.Soft (1, 0.929) vs Codon.Sto.Soft (2, 0.935) | 0.007 | 0.319 | No |
| Rank 1 vs 3 | Amino.Sto.Soft (1, 0.929) vs Amino.Det.Soft (3, 0.978) | 0.049 | 0.001 | Yes |
| Rank 1 vs 9 | Amino.Sto.Soft (1, 0.929) vs Lagrangian.Sto.Hard (9, 1.003) | 0.074 | 0.046 | Yes |

Paired t-tests across 12 proteins. Δ = difference in mean accessibility (kcal/mol).

The top two variants (Amino.Sto.Soft and Codon.Sto.Soft) are statistically equivalent (p = 0.319), forming an elite performance tier. Both employ stochastic soft discretization.

Accessibility-CAI

Table 6 summarizes pairwise statistical tests comparing top-ranked variants. Complete performance rankings for all 24 variants appear in Table S2.

**Table 6.** Pairwise Statistical Tests for Top Variant Rankings (Accessibility-CAI)

| Comparison | Variants (Rank, Mean kcal/mol) | Δ | p-value | Sig? |
|---|---|---|---|---|
| Rank 1 vs 2 | Amino.Sto.Hard +Penalty (1, 0.969) vs Codon.Sto.Hard (2, 0.977) | 0.008 | 0.427 | No |
| Rank 1 vs 3 | Amino.Sto.Hard +Penalty (1, 0.969) vs Codon.Det.Hard (3, 0.980) | 0.011 | 0.115 | No |
| Rank 1 vs 13 | Amino.Sto.Hard +Penalty (1, 0.969) vs C10 (13, 1.358) | 0.389 | <0.001 | Yes |

Paired t-tests across 12 proteins. Δ = difference in mean accessibility (kcal/mol).

The top three ID3 variants all employ hard discretization and achieve very similar performance (0.969–0.980 kcal/mol), unlike accessibility mode where soft variants dominate. All top 12 ID3 variants significantly outperform C10 combinatorial baseline (rank 13, 1.358 kcal/mol), demonstrating that full-sequence gradient-based optimization surpasses exhaustive local search even when constrained by CAI requirements.

##### S2.2.2. Constraint Mechanism Comparisons

The ID3 framework enforces amino acid conservation through three distinct mathematical approaches: *Codon Profile Constraint* applies soft constraints at the codon level based on synonymous codon probabilities; *Amino Matching Softmax* uses position-wise softmax to match target amino acid distributions; *Lagrangian Multiplier* enforces hard constraints through penalty terms. Despite these mathematical differences, we investigate whether these mechanisms achieve equivalent optimization performance.

Accessibility

Table 7 presents statistical comparison of the three constraint mechanisms (960 runs each: 48 configurations × 20 runs).

**Table 7.** Constraint Mechanism Statistical Comparison (Accessibility)

| Mechanism | Mean (kcal/mol) | Std | vs. Codon Diff | Cohen's d | p-value |
|---|---|---|---|---|---|
| Codon Profile | 0.972 | 0.195 | – | – | – |
| Amino Matching | 0.973 | 0.212 | -0.001 | -0.005 | 0.922 |
| Lagrangian | 1.231 | 0.349 | -0.259 | -0.918 | <0.001 |

Omnibus tests: ANOVA F=314.31 (p<0.001), Kruskal-Wallis H=420.44 (p<0.001)

Codon Profile and Amino Matching achieve statistically equivalent performance (p = 0.922), while Lagrangian underperforms by 26% (p<0.001, Cohen's d ≈ -0.9).

Accessibility-CAI

Table 8 presents statistical comparison combining both CAI integration methods (n = 1,920 per constraint).

Codon ≈ Amino equivalence persists in CAI mode (p = 0.602). Lagrangian's gap shrinks from 26% (accessibility) to 9–10% (CAI mode) but remains significant (p<0.001).

**Table 8.** Constraint Mechanism Statistical Comparison (Accessibility-CAI)

| Mechanism | Mean (kcal/mol) | Std | vs. Codon Diff | Cohen's d | p-value |
|---|---|---|---|---|---|
| Codon Profile | 1.266 | 0.357 | – | – | – |
| Amino Matching | 1.260 | 0.354 | -0.006 | -0.017 | 0.602 |
| Lagrangian | 1.386 | 0.437 | -0.120 | -0.30 | <0.001 |
| Combining 960 penalty-based + 960 no-penalty runs per mechanism | | | | | |

##### S2.2.3. Base Mode Comparisons

Accessibility

Table 9 compares the four base operational modes (720 runs each: 60 configurations × 20 runs).

**Table 9.** Base Mode Statistical Comparison (Accessibility)

| Base Mode | Mean (kcal/mol) | Std | Rank |
|---|---|---|---|
| Sto.Hard | 0.991 | 0.230 | 1 (best) |
| Det.Hard | 1.014 | 0.238 | 2 |
| Sto.Soft | 1.082 | 0.301 | 3 |
| Det.Soft | 1.148 | 0.344 | 4 (worst) |
| ANOVA: F=45.88, p<0.001 (significant differences among modes) | | | |

Stochastic hard mode achieves best performance (0.991 kcal/mol), with significant differences among all four modes (ANOVA F=45.88, p<0.001).

Accessibility-CAI

Table 10 compares the four base modes in CAI optimization (combining penalty + no-penalty runs).

**Table 10.** Base Mode Statistical Comparison (Accessibility-CAI)

| Base Mode | Mean (kcal/mol) | Std | Rank |
|---|---|---|---|
| Sto.Hard | 0.986 | 0.011 | 1 (best) |
| Det.Hard | 1.006 | 0.029 | 2 |
| Sto.Soft | 1.599 | 0.110 | 3 |
| Det.Soft | 1.624 | 0.111 | 4 (worst) |
| Combining penalty + no-penalty runs (6 variants per mode) | | | |

Hard modes significantly outperform soft modes. Stochastic vs. deterministic difference is marginal and not significant.

##### S2.2.4. Baseline Algorithm Comparisons

ID3 vs. Genetic Algorithm and Simulated Annealing

Genetic Algorithm (GA) and Simulated Annealing (SA) are widely-used stochastic optimization methods for sequence design. Both operate by iteratively proposing and evaluating sequence mutations without gradient information. Table 11 compares their performance against ID3 framework (2,880 normalized runs, accessibility). Detailed implementation parameters are provided below.

**Table 11.** ID3 Framework vs. Traditional Optimization Algorithms

| Method | Mean Accessibility (kcal/mol) | vs. ID3 (kcal/mol) | Effect Size (Cohen's d) | p-value |
|---|---|---|---|---|
| ID3 Framework | 1.030 | – | – | – |
| Genetic Algorithm | 2.090 | +1.060 (50% worse) | -2.74 | <0.001 |
| Simulated Annealing | 2.748 | +1.718 (62% worse) | -4.46 | <0.001 |
| Lower accessibility values indicate better performance. Effect sizes: very large. | | | | |

ID3 framework achieves 50–62% better accessibility than GA and SA ($p<0.001$, very large effect sizes $d < -2.7$), demonstrating substantial superiority of gradient-based optimization over stochastic search methods.

ID3 vs. C10 Combinatorial Optimal

The C10 baseline represents the theoretical optimum found by exhaustive combinatorial search over the first 10 codon positions (1,364,736 total combinations across 12 proteins, with search space ranging from 6,144 to 331,776 combinations per protein). Remaining positions were fixed to default codons.

95.6% of combinations satisfy CAI > 0.8, with only O15263 showing a measurable CAI constraint penalty (0.046 kcal/mol). This minimal average penalty (1.354 → 1.358 kcal/mol, +0.004) demonstrates that accessibility optimization and codon adaptation are largely compatible objectives.

**Accessibility-CAI Trade-off Landscape**: Figures 1 and 2 visualize the complete optimization landscape from 1,364,736 combinations. The scatter plot uses stratified sampling to show the relationship between CAI and accessibility, with red vertical line marking the CAI = 0.8 threshold. The distribution histograms show accessibility ranges for each protein across all combinations.

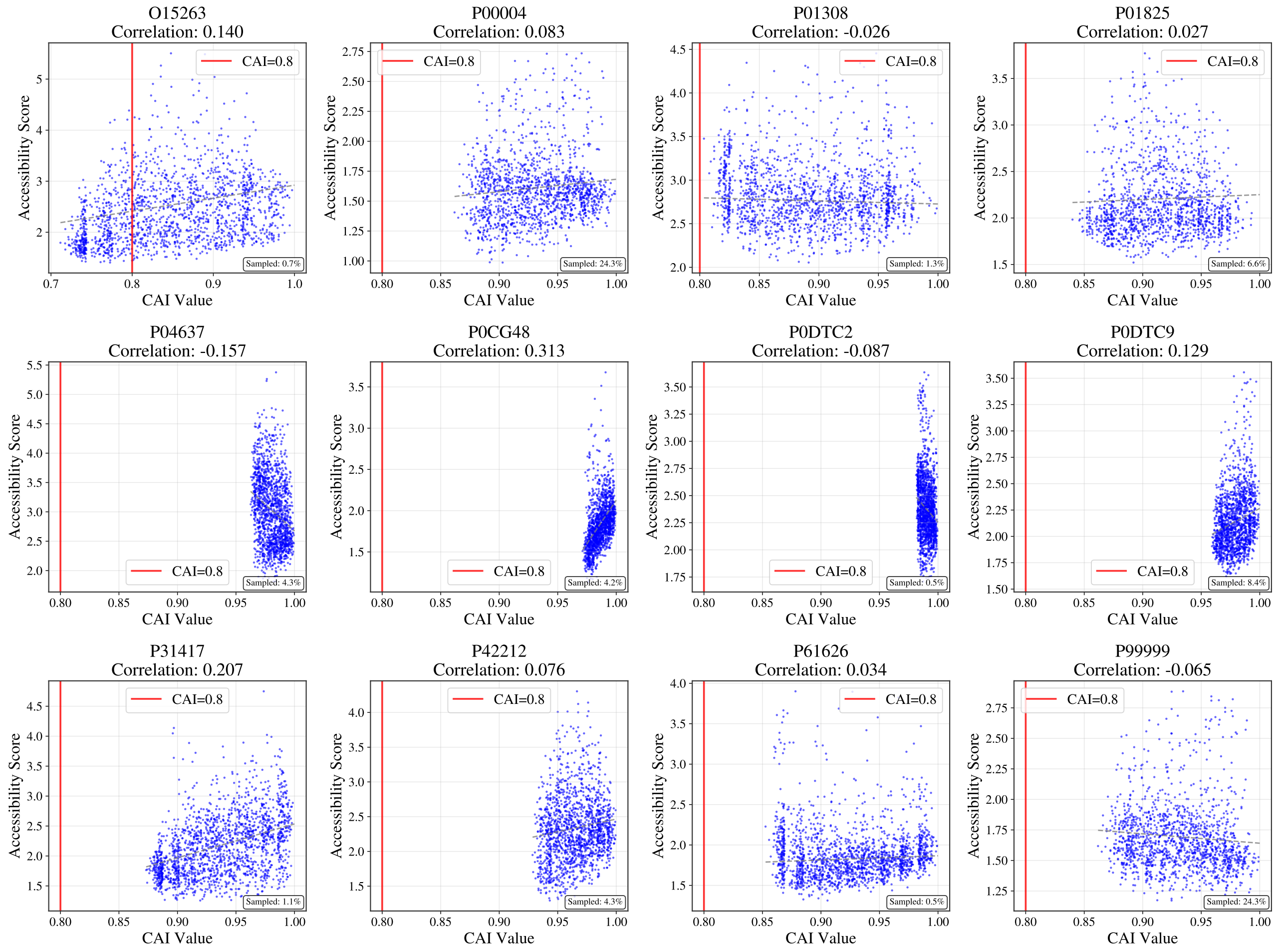


Fig. 1: Accessibility-CAI trade-off from C10 combinatorial exploration across 12 proteins. Each subplot shows a stratified sample representing the complete combinatorial space (total: 1,364,736 combinations). Red vertical line marks CAI = 0.8 threshold. Stratified sampling preserves distribution characteristics while maintaining visual clarity.

Performance Comparison

Table 12 compares C10 combinatorial baseline against top ID3 variants (paired t-tests across 12 proteins).

Full-Sequence vs. Partial-Window Optimization

These results demonstrate that full-sequence optimization with ID3 framework significantly outperforms even the theoretical combinatorial optimum for partial sequences. C10's 10-codon window achieves local theoretical optimum but misses global optimization potential. In contrast, ID3 optimizes all codon positions cooperatively, achieving 0.371 kcal/mol average improvement over C10 baseline. From a computational perspective, exhaustive search becomes prohibitive beyond 10–12 positions ($\sim 10^{15}$ combinations for full sequences), while gradient-based ID3 scales efficiently. Notably, CAI constraint (>0.8) adds minimal cost to C10 (1.354 → 1.358 kcal/mol, +0.004),

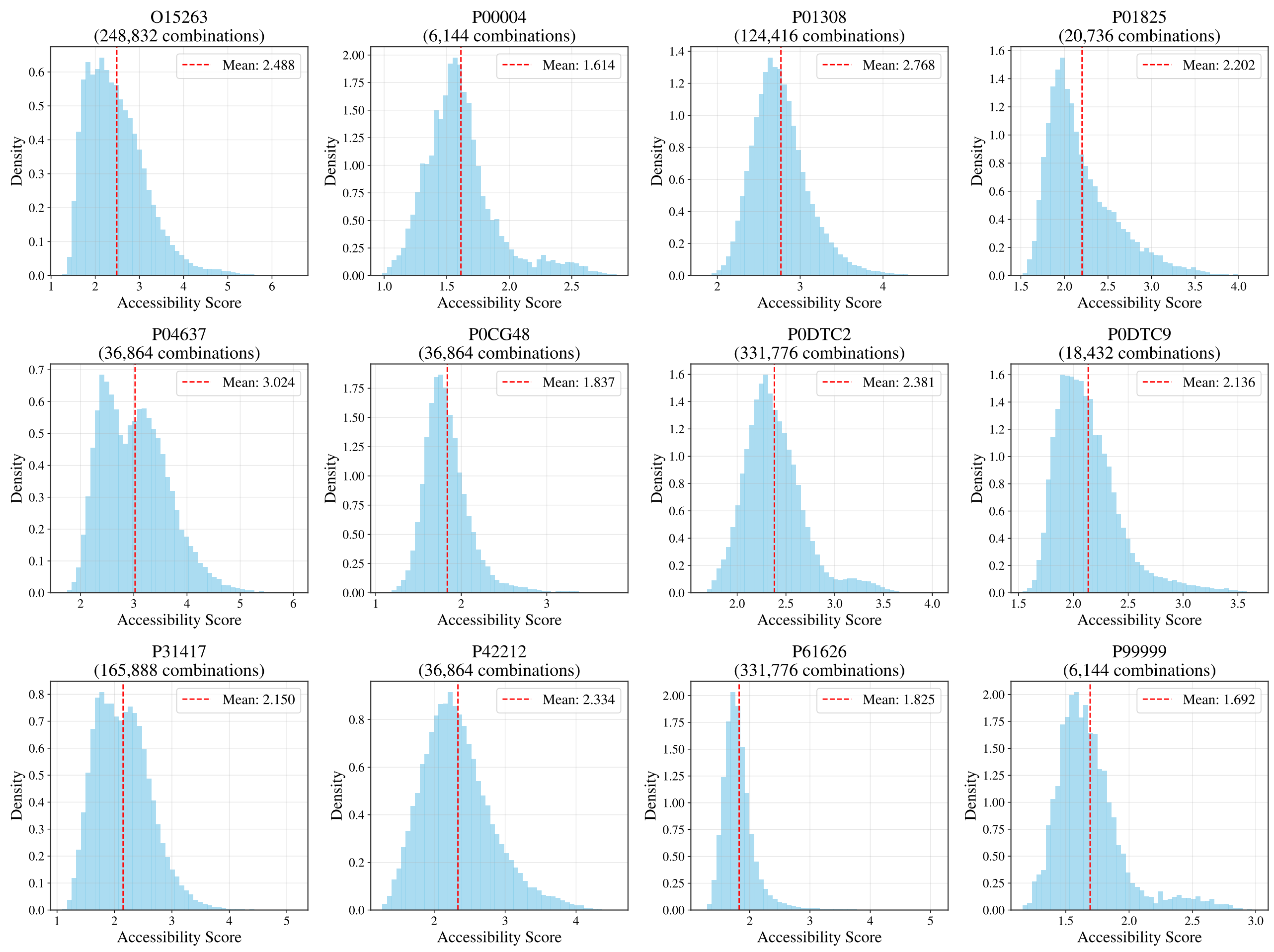


Fig. 2: Accessibility distributions from C10 combinatorial exploration across 12 proteins. Histograms show the complete accessibility landscape for all synonymous combinations of the first 10 codons (total: 1,364,736 combinations). Vertical lines indicate minimum accessibility values found for each protein.

**Table 12.** C10 Combinatorial Baseline vs. Top ID3 Variants

| Comparison | C10 Mean | ID3 Variant | ID3 Mean (kcal/mol) | Diff (kcal/mol) | t-stat | p-value |
|---|---|---|---|---|---|---|
| C10 vs. Rank 1 | 1.354 | Amino.Sto.Soft | 0.929 | +0.425 | +6.48 | <0.001 |
| C10 vs. Rank 2 | 1.354 | Codon.Sto.Soft | 0.935 | +0.419 | +6.36 | <0.001 |
| C10 vs. Top 10 | 1.354 | Average of top 10 | 0.983 | +0.371 | – | <0.01 |
| C10 (CAI > 0.8) | 1.358 | Rank 13 in Table S2 (outperformed by all top 12 ID3 variants) | | | | |

indicating most optimal combinations naturally satisfy codon adaptation requirements. This comparison validates the necessity of full-sequence gradient-based optimization and establishes C10 as a strong theoretical baseline that ID3 consistently surpasses.

##### S2.2.5. CAI Integration Methods: Penalty vs. No-Penalty

Two approaches integrate CAI constraints: penalty-based (multi-objective joint loss) and no-penalty (CAI-aware discretization). Table 13 compares their performance across all 12 base variants (n = 2,880 each).

Both CAI integration methods achieve statistically equivalent performance (difference = 0.005 kcal/mol, p = 0.661), with equivalence persisting across all constraint types and base modes. Detailed ablation study and methodological comparison in Section S8.

##### S2.2.6. Summary of Statistical Findings

1. **Constraint Mechanism Equivalence**: Codon Profile and Amino Matching show statistically equivalent performance (difference ¡ 0.1%, $p = 0.922$, Cohen's $d = -0.005$), indicating these mechanisms are functionally interchangeable for accessibility optimization.

**Table 13.** Penalty vs. No-Penalty CAI Integration Comparison

| Method | Mean (kcal/mol) | Std | Cohen's d | p-value |
|---|---|---|---|---|
| Penalty-based | 1.306 | 0.387 | 0.012 | 0.661 |
| No-Penalty | 1.301 | 0.390 | | |
| **By Constraint**: Codon Δ=+0.008, Amino Δ=-0.006, Lagrangian Δ=+0.011 (all p>0.05) | | | | |
| **By Base Mode**: Det.Soft Δ=-0.008, Det.Hard Δ=+0.012, Sto.Soft Δ=+0.008, Sto.Hard Δ=+0.004 (all p>0.05) | | | | |

2. **Lagrangian Underperformance**: Lagrangian Multiplier performs significantly worse than both Codon and Amino mechanisms (0.258-0.259 kcal/mol worse representing 26% degradation, large effect size $d \approx -0.9$, $p < 0.001$).
3. **Top Variant Similarity**: Rank 1 (Amino.Sto.Soft, 0.929 kcal/mol) and Rank 2 (Codon.Sto.Soft, 0.935 kcal/mol) are statistically indistinguishable ($p = 0.319$), with performance difference of only 0.7%. Significant differences emerge starting from Rank 3 ($p = 0.001$).
4. **Base Mode Performance**: Stochastic hard mode achieves best overall performance (0.991 kcal/mol) compared to other modes (Det.Hard: 1.014, Sto.Soft: 1.082, Det.Soft: 1.148), with significant differences confirmed by ANOVA ($F = 45.88$, $p < 0.001$).
5. **Framework Superiority over Baselines**: ID3 framework provides substantial improvements over traditional optimization methods: 50.0% better than Genetic Algorithm (1.060 kcal/mol improvement, $d = -2.74$) and 61.9% better than Simulated Annealing (1.718 kcal/mol improvement, $d = -4.46$), both highly significant ($p < 0.001$).
6. **Full-Sequence Optimization Advantage**: All top 10 ID3 variants significantly outperform C10 combinatorial baseline (average 0.370 kcal/mol better, all $p < 0.01$), demonstrating that full-sequence gradient-based optimization surpasses exhaustive search on partial sequences.

#### S3. Notation and Symbols

This section provides a comprehensive reference for all mathematical symbols used throughout the main paper and supplementary material, organized by functional categories.

**Table 14.** ID3 Framework Core Symbols

| Symbol | Definition | Dimension |
|---|---|---|
| $\Theta$ | Learnable parameters (dimension varies by constraint) | See constraint-specific definitions |
| $\Theta_{\text{codon}}$ | Codon-level parameters (Codon Profile) | $\mathbb{R}^{\sum_j \lvert\mathcal{C}(y_j)\rvert}$ |
| $\mathbf{P}$ | RNA probability distribution (sequence profile) | $[0,1]^{L\times 4}$ |
| $\mathbf{S}$ | RNA sequence (discrete or soft) | $\{A,U,G,C\}^L$ or $[0,1]^{L\times 4}$ |
| $\mathbf{P}_{\text{codon}}$ | Codon probability distribution | $[0,1]^M$ |
| $T$ | Unified transformation function | $\Theta \mapsto \mathbf{S}$ |
| $\Pi$ | Parameter-to-probability transformation | $\Theta \mapsto \mathbf{P}$ |
| $\Psi$ | Probability-to-sequence transformation | $\mathbf{P} \mapsto \mathbf{S}$ |
| $\sigma$ | Temperature parameter | $\mathbb{R}^+$ |
| $T^{\text{det.soft}}(\Theta)$ | Deterministic soft transformation | $\Theta \mapsto \mathbf{S}$ |
| $T^{\text{det.hard}}(\Theta)$ | Deterministic hard transformation | $\Theta \mapsto \mathbf{S}$ |
| $T^{\text{sto.soft}}(\Theta)$ | Stochastic soft transformation | $\Theta \mapsto \mathbf{S}$ |
| $T^{\text{sto.hard}}(\Theta)$ | Stochastic hard transformation | $\Theta \mapsto \mathbf{S}$ |
| $T^{\text{soft}}$ | Generic soft transformation | $\mathbf{P} \mapsto \mathbf{S}$ |
| $T^{\text{hard}}$ | Generic hard transformation | $\mathbf{P} \mapsto \mathbf{S}$ |

**Table 15.** Biological Sequence Symbols

| Symbol | Definition | Dimension |
|---|---|---|
| $\mathbf{y}$ | Target amino acid sequence | $\{\text{AA}\}^M$ |
| $M$ | Number of amino acids (sequence length) | $\mathbb{N}$ |
| $L$ | RNA sequence length ($L = 3M$) | $\mathbb{N}$ |
| $\mathcal{C}(y_j)$ | Set of valid codons for amino acid $y_j$ | $\{\text{codon}\}$ |
| $\mathcal{E}[c]$ | One-hot encoding of codon $c$ | $\{0,1\}^{3\times4}$ |
| $g$ | Gumbel noise | $\mathbb{R}$ |
| $\theta_{j,c}$ | Parameter for codon $c$ at position $j$ (Codon Profile) | $\mathbb{R}$ |
| $\Theta^{(j)}$ | RNA parameters for position $j$ (Amino Matching) | $\mathbb{R}^{3\times4}$ |

**Table 16.** Application-Specific Symbols

| Symbol | Definition | Dimension |
|---|---|---|
| $f_{\text{model}}$ | Pre-trained predictor (DeepRaccess) | $\mathbb{R}^L \mapsto \mathbb{R}$ |
| $f_{\text{access}}$ | RNA accessibility prediction function | $\mathbb{R}^L \mapsto \mathbb{R}$ |
| $\mathcal{L}_{\text{access}}$ | Accessibility loss function | $\mathbb{R}$ |
| CAI | Codon Adaptation Index | $[0,1]$ |
| ECAI | Expected CAI | $[0,1]$ |
| $w_c$ | CAI weight for codon $c$ (see Table S2_additional) | $[0,1]$ |
| $\gamma$ | Interpolation parameter (binary search) | $[0,1]$ |
| threshold | Target CAI threshold | $[0,1]$ |
| $\Pi_{\text{codon}}$ | Codon-specific probability function (Codon Profile) | $\Theta_{\text{codon}} \mapsto \mathbf{P}_{\text{codon}}$ |
| $\Pi_{\text{amino}}$ | Amino-based probability function (Amino Matching) | $\Theta \mapsto \mathbf{P}_{\text{codon}}$ |
| $\text{Onehot}(\cdot)$ | Vectorized one-hot encoding function | $\mathbf{P}_{\text{codon}} \mapsto \mathbf{S}$ |
| $R_{\text{nuc}}(\cdot)$ | Vectorized nucleotide reconstruction function | $\mathbf{P}_{\text{codon}} \mapsto \mathbf{S}$ |
| $\text{soft\_embedding}^{(i)}$ | Soft embedding at position $i$ | $\mathbb{R}^d$ |
| $\text{embedding}[\cdot]$ | Trained embedding layer | $\mathbb{R}^d$ |
| $5\text{'UTR}_{70}$ | 5' Untranslated Region (70 nt) | RNA sequence |
| $3\text{'UTR}_{63}$ | 3' Untranslated Region (63 nt) | RNA sequence |
| $\text{ATG} - 19 : +15$ | Ribosome binding region for accessibility optimization | RNA region |
| $\text{CDS}_{\text{var}}$ | Coding Sequence (variable length) | RNA sequence |

**Table 17.** Optimization and Convergence Symbols

| Symbol | Definition | Dimension |
|---|---|---|
| $\mathcal{L}$ | General loss function | $\mathbb{R}$ |
| $F(\Theta)$ | Objective function for optimization | $\mathbb{R}$ |
| $\nabla$ | Gradient operator | Vector field |
| $\eta$ | Learning rate | $\mathbb{R}^+$ |
| $\lambda$ | Lagrangian multiplier | $\mathbb{R}^+$ |
| $\mathcal{C}(\mathbf{P})$ | Constraint penalty function | $\mathbb{R}^+$ |
| $L_{\text{composite}}$ | Composite Lipschitz constant | $\mathbb{R}^+$ |
| $L_{\mathcal{L}}$ | Loss function Lipschitz constant | $\mathbb{R}^+$ |
| $B_{\text{STE}}$ | STE bias bound | $\mathbb{R}^+$ |
| $\Delta_{\max}$ | Maximum probability difference | $[0,1]$ |
| $\epsilon$ | Convergence tolerance | $\mathbb{R}^+$ |
| $k$ | Top-k position selection parameter | $\mathbb{N}$ |
| $\text{potential}_i$ | Improvement potential at position $i$ | $[0,1]$ |
| score | Composite evaluation score | $[0,1]$ |
| $h(\cdot)$ | Sequence hash function (duplicate detection) | $\mathbb{N} \mapsto$ String |
| $\alpha_{\text{CAI}}, \alpha_{\text{prob}}$ | Score weighting parameters | $[0,1]$ |

**Table 18.** ID3 Variant Notation (V40 Descriptive Naming)

| Notation | Description |
|---|---|
| **Base Operational Modes** | |
| Det.Soft | Deterministic parameters with soft (continuous) output |
| Det.Hard | Deterministic parameters with hard (discrete) output |
| Sto.Soft | Stochastic parameters with soft (continuous) output |
| Sto.Hard | Stochastic parameters with hard (discrete) output |
| **Constraint Mechanisms (ordered by presentation in text)** | |
| Codon | Codon Profile Constraint variants |
| Amino | Amino Matching Softmax variants |
| Lagrangian | Lagrangian Multiplier variants |
| **Complete Variant Naming** | |
| Codon.Det.Soft | Codon Profile Constraint + Deterministic Soft |
| Codon.Det.Hard | Codon Profile Constraint + Deterministic Hard |
| Codon.Sto.Soft | Codon Profile Constraint + Stochastic Soft |
| Codon.Sto.Hard | Codon Profile Constraint + Stochastic Hard |
| Amino.Det.Soft | Amino Matching Softmax + Deterministic Soft |
| Amino.Det.Hard | Amino Matching Softmax + Deterministic Hard |
| Amino.Sto.Soft | Amino Matching Softmax + Stochastic Soft |
| Amino.Sto.Hard | Amino Matching Softmax + Stochastic Hard |
| Lagrangian.Det.Soft | Lagrangian Multiplier + Deterministic Soft |
| Lagrangian.Det.Hard | Lagrangian Multiplier + Deterministic Hard |
| Lagrangian.Sto.Soft | Lagrangian Multiplier + Stochastic Soft |
| Lagrangian.Sto.Hard | Lagrangian Multiplier + Stochastic Hard |

**Table 19.** Additional Symbols for Convergence Analysis

| Symbol | Definition | Dimension |
|---|---|---|
| *Probability and Distribution Symbols* | | |
| $p_k^{(i)}$ | Probability of nucleotide $k \in \{A, U, G, C\}$ at position $i$ | $[0, 1]$ |
| $\mathbf{P}_{\text{codon}}$ | Codon probability vector | $[0, 1]^M$ |
| $P_{\text{codon}_{j,c}}$ | Probability of codon $c$ at amino acid position $j$ | $[0, 1]$ |
| $\mathbf{P}^{(j)}$ | Probability matrix for codon at position $j$ | $[0, 1]^{3\times4}$ |
| *Optimization Bounds and Constants* | | |
| $\epsilon_{\min}$ | Minimum probability bound (numerical stability) | $\mathbb{R}^+$ |
| $\sigma_{\min}, \sigma_{\max}$ | Temperature parameter bounds | $\mathbb{R}^+$ |
| $C_f$ | Lipschitz constant for function $f_{\text{model}}$ | $\mathbb{R}^+$ |
| $C_{\mathcal{L}}$ | Lipschitz constant for loss function $\mathcal{L}$ | $\mathbb{R}^+$ |
| $C_{\text{composite}}$ | Composite Lipschitz constant | $\mathbb{R}^+$ |
| $B$ | Upper bound for gradient magnitude | $\mathbb{R}^+$ |
| $B_{\text{STE}}$ | STE gradient bias bound: $\|g_{\text{STE}} - \nabla f\| \leq B_{\text{STE}}$ | $\mathbb{R}^+$ |
| $C_{\text{STE}}$ | STE constant: $C_{\text{STE}} = C_{\mathcal{L}} \cdot C_f$ | $\mathbb{R}^+$ |
| $\Delta_{\max}$ | Maximum probability difference in distribution | $[0, 1]$ |
| *Iteration and Convergence Symbols* | | |
| $t$ | Iteration index | $\mathbb{N}$ |
| $N$ | Total number of iterations | $\mathbb{N}$ |
| $\Theta^{(t)}$ | Parameters at iteration $t$ | $\mathbb{R}^n$ |
| $\Theta^*$ | Optimal parameters at convergence | $\mathbb{R}^n$ |
| *Stochastic Optimization Symbols* | | |
| $\sigma_G$ | Standard deviation of Gumbel noise | $\mathbb{R}^+$ |
| $\xi^{(t)}$ | Stochastic noise contribution to gradient at iteration $t$ | $\mathbb{R}^n$ |
| $\nabla F_{\text{det}}(\Theta)$ | Deterministic gradient component | $\mathbb{R}^n$ |
| *Constraint Optimization Symbols* | | |
| $\mu$ | Strong convexity constant | $\mathbb{R}^+$ |
| $g^{(t)}$ | Subgradient of constraint at iteration $t$ (Lagrangian) | $\mathbb{R}$ |
| $\eta_0$ | Initial learning rate | $\mathbb{R}^+$ |
| $\eta_t$ | Learning rate at iteration $t$ | $\mathbb{R}^+$ |
| $D_{\text{feas}}$ | Diameter of feasible region | $\mathbb{R}^+$ |
| *Mathematical Operators* | | |
| $\delta_{ij}$ | Kronecker delta: 1 if $i = j$, 0 otherwise | $\{0, 1\}$ |
| $\text{vec}(\mathbf{A})$ | Vectorization: stack columns of matrix $\mathbf{A}$ | Function |
| $\langle \mathbf{a}, \mathbf{b} \rangle$ | Inner product of vectors $\mathbf{a}$ and $\mathbf{b}$ | $\mathbb{R}$ |
| *CAI Optimization Symbols* | | |
| $\mathbf{w}$ | CAI-optimal codon probability distribution | $[0, 1]^M$ |
| $w_c$ | CAI weight for codon $c$ (relative adaptiveness) | $[0, 1]$ |
| $\mathcal{S}_{\text{valid}}$ | Valid amino-acid-preserving sequence space | Set |
| $\mathbf{v}_c$ | Feature vector for codon $c$ (Amino Matching) | $\mathbb{R}^{12}$ |
| $X_{ij}$ | Count of codon $j$ for amino acid $i$ in reference | $\mathbb{N}$ |
| $n_i$ | Number of synonymous codons for amino acid $i$ | $\mathbb{N}$ |

#### S4. Mathematical Foundations and Assumptions

##### S4.1. Fundamental Assumptions

The convergence analysis of the ID3 framework relies on three fundamental assumptions that reflect the practical characteristics of pre-trained neural networks and RNA sequence optimization.

**Assumption 1** (Pre-trained Model Local Properties) The pre-trained model $f_{\text{model}}$ exhibits local regularity in the optimization region. Here, $\mathbf{x} = T(\Theta)$ represents the model input (the RNA sequence representation obtained from parameters $\Theta$ via transformation $T$).

- **Local Lipschitz continuity** [Nesterov, 2004]: There exists a constant $C_f > 0$ such that for any two inputs $\mathbf{x}_1, \mathbf{x}_2$ encountered during optimization:
$$|f_{\text{model}}(\mathbf{x}_1) - f_{\text{model}}(\mathbf{x}_2)| \leq C_f \|\mathbf{x}_1 - \mathbf{x}_2\| \quad (1)$$
- **Gradient boundedness**: The gradient magnitude is bounded: $\|\nabla f_{\text{model}}(\mathbf{x})\| \leq B < \infty$ for finite constant $B$.

**Rationale**: Well-trained neural networks typically exhibit smooth behavior in regions containing biologically plausible sequences. The pre-trained DeepRaccess model, trained on diverse RNA sequences, maintains regularity within the manifold of functional RNA sequences encountered during optimization.

**Assumption 2** (Optimization Trajectory Boundedness) The probability distributions and temperature parameters remain within practical bounds during optimization. Here, $p_k^{(i)}$ denotes the probability of nucleotide $k \in \{A, U, G, C\}$ at position $i$ in the RNA sequence, and $\sigma$ is the temperature parameter controlling the softness of the distribution.

- **Probability bounds**: All nucleotide probabilities satisfy $p_k^{(i)} \in [\epsilon_{\min}, 1 - \epsilon_{\min}]$ where $\epsilon_{\min} > 0$ prevents probability collapse (i.e., ensures no probability approaches exactly 0 or 1).
- **Temperature bounds**: The temperature parameter remains bounded: $\sigma \in [\sigma_{\min}, \sigma_{\max}]$ with $0 < \sigma_{\min} \leq \sigma_{\max} < \infty$.

**Practical significance**: These bounds prevent numerical instabilities and ensure that the softmax transformation maintains meaningful probability distributions. In practice, $\epsilon_{\min} = 10^{-6}$ and $\sigma \in [0.1, 2.0]$ provide stable optimization behavior while preserving biological interpretability.

**Assumption 3** (Loss Function Regularity) The loss function $\mathcal{L}$ possesses basic regularity properties:

- **Continuous differentiability**: $\mathcal{L}$ is continuously differentiable with well-defined gradients.
- **Lipschitz continuous gradients**: There exists a constant $C_{\mathcal{L}} > 0$ such that:
$$\|\nabla \mathcal{L}(z_1) - \nabla \mathcal{L}(z_2)\| \leq C_{\mathcal{L}} \|z_1 - z_2\| \quad (2)$$
where $z_1, z_2$ are generic function inputs.

**Note**: Standard loss functions such as MSE, cross-entropy, and smooth approximations of ranking losses naturally satisfy these conditions.

##### S4.2. Optimization Objective

The general ID3 optimization problem takes the form:
$$\min_{\Theta} f_{\text{model}}(T(\Theta)) \quad (3)$$

where $f_{\text{model}}$ is the fixed pre-trained model, $T(\Theta) = \Psi(\Pi(\theta))$ is the unified transformation function following the three-layer hierarchical architecture (Parameter Level → Probability Level → Output Level), and $\Theta$ represents optimizable parameters.

The objective function represents a composition of three functions:
$$F(\Theta) = \mathcal{L}(f_{\text{model}}(T(\Theta))) = (\mathcal{L} \circ f_{\text{model}} \circ T)(\Theta) \quad (4)$$

#### S5. Convergence Analysis

##### S5.1. Base Variant Convergence Analysis

###### S5.1.1. Det.Soft: Deterministic Soft Convergence

Det.Soft represents the most theoretically tractable variant, using deterministic softmax transformations without additional complexity.

**Theorem 1** (Det.Soft Convergence) *Under Assumptions 1-3, gradient descent for Det.Soft mode with learning rate $\eta < 2/C_{composite}$ satisfies:*
$$\lim_{t \to \infty} \|\nabla F(\Theta^{(t)})\| = 0 \quad (5)$$
*where $t$ denotes the iteration index, $\Theta^{(t)}$ represents the parameters at iteration $t$, and the composite Lipschitz constant is:*
$$C_{composite} = C_{\mathcal{L}} \cdot C_f \cdot \frac{1}{\sigma} \quad (6)$$

*Proof* We establish convergence through a three-step analysis leveraging the composition structure $F = \mathcal{L} \circ f_{\text{model}} \circ T$.

*Step 1: Lipschitz Constant Analysis.* The composite Lipschitz constant is determined by the chain rule. For the gradient of the composite function:

$$\|\nabla F(\Theta)\| = \|\nabla \mathcal{L}(f_{\text{model}}(T(\Theta))) \cdot \nabla f_{\text{model}}(T(\Theta)) \cdot \nabla T(\Theta)\| \tag{7}$$

For the softmax transformation $T(\Theta)$, the gradient with respect to parameters has magnitude bounded by $\|\nabla T(\Theta)\| \leq \frac{1}{\sigma}$. This follows from the softmax gradient formula. Recall that $\theta_j^{(i)}$ denotes the parameter (logit) for nucleotide $j$ at position $i$, and $p_k^{(i)}$ is the corresponding probability. The gradient of probability with respect to the parameter is:

$$\frac{\partial p_k^{(i)}}{\partial \theta_j^{(i)}} = \frac{1}{\sigma}[p_k^{(i)}(\delta_{kj} - p_j^{(i)})] \tag{8}$$

where $\delta_{kj}$ is the Kronecker delta function ($\delta_{kj} = 1$ if $k = j$, and $\delta_{kj} = 0$ otherwise). The bound follows from $|p_k^{(i)}(\delta_{kj} - p_j^{(i)})| \leq p_k^{(i)} \leq 1$.

Combining with Assumptions 1 and 3, the composite Lipschitz constant becomes:

$$C_{\text{composite}} = C_{\mathcal{L}} \cdot C_f \cdot \frac{1}{\sigma} \tag{9}$$

*Step 2: Descent Lemma Application.* For the $C_{\text{composite}}$-smooth function $F$, the standard descent lemma gives:

$$F(\Theta^{(t+1)}) \leq F(\Theta^{(t)}) - \frac{\eta}{2}\|\nabla F(\Theta^{(t)})\|^2 \tag{10}$$

provided $\eta < 2/C_{\text{composite}}$. This inequality holds for gradient descent updates $\Theta^{(t+1)} = \Theta^{(t)} - \eta \nabla F(\Theta^{(t)})$.

*Step 3: Telescoping Analysis and Convergence.* Summing the descent inequality over iterations $t = 0, \ldots, N-1$:

$$\sum_{t=0}^{N-1} \|\nabla F(\Theta^{(t)})\|^2 \leq \frac{2}{\eta}[F(\Theta^{(0)}) - F(\Theta^{(N)})] \tag{11}$$

Since $F$ is bounded below (the loss function $\mathcal{L}$ has a lower bound by assumption) and $F(\Theta^{(0)})$ is finite, the right-hand side is bounded. Therefore, $\sum_{t=0}^{\infty} \|\nabla F(\Theta^{(t)})\|^2 < \infty$, which implies:

$$\lim_{t\to\infty} \|\nabla F(\Theta^{(t)})\| = 0 \tag{12}$$

□

##### S5.1.2. Sto.Soft: Stochastic Soft Convergence

Sto.Soft extends Det.Soft by adding Gumbel noise [Jang et al., 2016] for exploration, introducing stochastic elements while maintaining the core convergence structure.

**Theorem 2** (Sto.Soft Stochastic Convergence) *With Gumbel noise, Sto.Soft mode satisfies:*

$$E[\|\nabla F(\Theta^{(t)})\|^2] \to 0 \textit{ as } t \to \infty \tag{13}$$

*Proof* We analyze stochastic convergence through gradient decomposition and expectation bounds.

*Step 1: Gradient Decomposition.* The stochastic gradient can be decomposed as:

$$\nabla F(\Theta^{(t)}) = \nabla F_{\text{det}}(\Theta^{(t)}) + \xi^{(t)} \tag{14}$$

where $\nabla F_{\text{det}}(\Theta^{(t)})$ is the deterministic gradient (without noise), and $\xi^{(t)}$ represents the stochastic noise contribution to the gradient at iteration $t$ from Gumbel sampling. This noise is zero-mean ($E[\xi^{(t)}] = 0$) with bounded variance $E[\|\xi^{(t)}\|^2] \leq \sigma_G^2$, where $\sigma_G$ denotes the standard deviation of the Gumbel noise.

*Step 2: Expected Descent Inequality.* Applying the descent lemma to the expected objective and using the bias-variance decomposition:

$$\begin{aligned} E[F(\Theta^{(t+1)})] &\leq E[F(\Theta^{(t)})] - \eta E[\langle \nabla F(\Theta^{(t)}), \nabla F_{\text{det}}(\Theta^{(t)})\rangle] \\ &\quad + \frac{\eta^2 C_{\text{composite}}}{2} E[\|\nabla F(\Theta^{(t)})\|^2] \\ &= E[F(\Theta^{(t)})] - \eta E[\|\nabla F_{\text{det}}(\Theta^{(t)})\|^2] \\ &\quad + \frac{\eta^2 C_{\text{composite}}}{2}(E[\|\nabla F_{\text{det}}(\Theta^{(t)})\|^2] + \sigma_G^2) \end{aligned} \tag{15}$$

where we used $E[\langle \nabla F_{\text{det}}(\Theta^{(t)}), \xi^{(t)}\rangle] = 0$ due to independence.

*Step 3: Convergence in Expectation.* Rearranging the inequality:

$$E[\|\nabla F_{\text{det}}(\Theta^{(t)})\|^2] \leq \frac{2}{\eta(2-\eta C_{\text{composite}})}[E[F(\Theta^{(t)})] - E[F(\Theta^{(t+1)})]] + \frac{\eta C_{\text{composite}}\sigma_G^2}{2-\eta C_{\text{composite}}} \tag{16}$$

For learning rate $\eta < 2/C_{\text{composite}}$, telescoping over iterations and using boundedness of $F$ yields:

$$\sum_{t=0}^{\infty} E[\|\nabla F_{\text{det}}(\Theta^{(t)})\|^2] < \infty \tag{17}$$

Therefore, $E[\|\nabla F(\Theta^{(t)})\|^2] \to 0$ as $t \to \infty$. □

**S5.1.3. Det.Hard and Sto.Hard: Straight-Through Estimator Variants**

The Straight-Through Estimator (STE) [Bengio et al., 2013] is a gradient approximation technique used when the forward pass involves discrete operations (e.g., argmax, sampling) that are non-differentiable. In the ID3 framework:

- **Forward pass**: Use discrete/hard sequences (e.g., via argmax or Gumbel sampling with discretization)
- **Backward pass**: Compute gradients as if the forward pass used soft probabilities (ignoring the discretization)

This approximation introduces a gradient bias $B_{\text{STE}}$, which affects convergence properties.

**Theorem 3** (Straight-Through Estimator Convergence) *For any differentiable optimization problem with Lipschitz-continuous objective function $f : \mathbb{R}^n \to \mathbb{R}$ with Lipschitz constant $L_f$, when the Straight-Through Estimator (STE) is applied with gradient estimation bias bounded by $B_{STE}$, the optimization converges to an $O(B_{STE})$-neighborhood of the true optimum rather than exact convergence.*

*Proof* Let $g_{\text{STE}}(x)$ denote the biased gradient estimate produced by STE (which uses gradients from the soft distribution while discretizing the forward pass), and let $\nabla f(x)$ denote the true gradient. The STE bias satisfies $\|g_{\text{STE}}(x) - \nabla f(x)\| \leq B_{\text{STE}}$ for all $x$.

Consider the gradient descent update $x^{(t+1)} = x^{(t)} - \eta g_{\text{STE}}(x^{(t)})$. For a stationary point $x^*$ of the true objective, we have:

$$\begin{aligned} \|x^{(t+1)} - x^*\| &= \|x^{(t)} - \eta g_{\text{STE}}(x^{(t)}) - x^*\| \\ &= \|x^{(t)} - x^* - \eta \nabla f(x^{(t)}) - \eta(g_{\text{STE}}(x^{(t)}) - \nabla f(x^{(t)}))\| \end{aligned} \tag{18}$$

Using the descent property of gradient descent and the bias bound:

$$\|x^{(t+1)} - x^*\| \leq (1-\eta\mu)\|x^{(t)} - x^*\| + \eta B_{\text{STE}} \tag{19}$$

where $\mu > 0$ is the strong convexity constant (a measure of the curvature of the objective function near the optimum).

Taking the limit, the algorithm converges to a neighborhood of radius $\frac{\eta B_{\text{STE}}}{\eta\mu} = \frac{B_{\text{STE}}}{\mu} = O(B_{\text{STE}})$ around the true optimum. □

This fundamental result leads to the following extensions:

**Corollary 4** (Det.Hard Convergence) *Det.Hard (STE + Deterministic) satisfies:*

$$\textit{Det.Soft: } \|\nabla F(\Theta^*)\| = 0 \Rightarrow \textit{Det.Hard: } \|\nabla F(\Theta^*)\| \leq B_{STE} \tag{20}$$

**Corollary 5** (Sto.Hard Convergence) *Sto.Hard (STE + Stochastic) satisfies:*

$$E[\|\nabla F(\Theta^*)\|] \leq \sqrt{B_{STE}^2 + \sigma_G^2} \tag{21}$$

The STE bias bound quantifies the error introduced by using soft gradients while discretizing the forward pass. It is given by:

$$B_{\text{STE}} \leq \frac{C_{\text{STE}}}{\sigma} \cdot \Delta_{\max} \tag{22}$$

where:

- $B_{\text{STE}}$ is the maximum difference between the biased STE gradient and the true gradient: $\|g_{\text{STE}}(x) - \nabla f(x)\| \leq B_{\text{STE}}$
- $C_{\text{STE}}$ is a constant depending on the Lipschitz constants of the model $f_{\text{model}}$ and loss function $\mathcal{L}$ (specifically, $C_{\text{STE}} = C_{\mathcal{L}} \cdot C_f$)
- $\Delta_{\max} = \max_{i,k,k'} |p_k^{(i)} - p_{k'}^{(i)}|$ is the maximum probability difference between any two nucleotides at any position in the softmax distribution
- $\sigma$ is the temperature parameter (lower temperature increases sharpness and thus increases bias)

The STE bias arises from the discrepancy between computing gradients using the soft probability distribution and evaluating losses using discrete (hard) sequences. When the temperature $\sigma$ is small (sharper distribution), the soft and hard sequences are more similar, reducing the bias. The bound shows that STE bias decreases as $1/\sigma$ when probabilities become more concentrated.

#### S5.2. Constraint Mechanism Convergence Analysis

##### S5.2.1. Vectorized Transformation Functions

Before presenting constraint-specific convergence analysis, we define the vectorized transformation functions introduced in the main paper that convert codon probabilities to sequence representations.

**Hard Mode Discretization** (main paper Equation 7):

$$\text{Onehot}(\mathbf{P}_{\text{codon}}) = \{\mathcal{E}[\arg\max_{c\in\mathcal{C}(y_j)} P_{\text{codon}_{j,c}}]\}_{j=1}^{M} \tag{23}$$

This function selects the highest-probability codon at each position $j$ and returns its one-hot encoding $\mathcal{E}[c]$, producing a discrete RNA sequence.

**Soft Mode Reconstruction** (main paper Equation 8):

$$R_{\text{nuc}}(\mathbf{P}_{\text{codon}}) = \{\sum_{c\in\mathcal{C}(y_j)} P_{\text{codon}_{j,c}} \cdot \mathcal{E}[c]\}_{j=1}^{M} \tag{24}$$

This function reconstructs nucleotide probabilities as weighted sums of codon encodings, producing a continuous probability distribution over nucleotides while preserving amino acid constraints.

**Transformation Patterns by Constraint Type** (main paper Table 2):

- **Codon Profile Constraint & Amino Matching Softmax:** Both use:
  - Soft mode: $T^{\text{soft}} = R_{\text{nuc}}(\mathbf{P}_{\text{codon}})$
  - Hard mode: $T^{\text{hard}} = \text{Onehot}(\mathbf{P}_{\text{codon}})$

  They differ in how $\mathbf{P}_{\text{codon}}$ is computed: via $\Pi_{\text{codon}}(\Theta_{\text{codon}})$ for Codon Profile versus $\Pi_{\text{amino}}(\Theta^{(j)})$ for Amino Matching.
- **Lagrangian Multiplier:** Uses:
  - Soft mode: $T^{\text{soft}} = \mathbf{P}$ (raw nucleotide probabilities)
  - Hard mode: $T^{\text{hard}} = \text{Onehot}(\Pi_{\text{amino}}(\mathbf{P}))$ (projection then discretization)

These unified transformation functions enable consistent analysis across all constraint mechanisms while respecting their distinct approaches to amino acid preservation.

##### S5.2.2. Codon Profile Convergence

Codon Profile operates in a reduced parameter space where amino acid sequences are guaranteed to be preserved throughout optimization.

**Parameter Space** (main paper Equation 1): $\Theta_{\text{codon}} = \{\theta_{j,c} : c \in \mathcal{C}(y_j), j \in [1, M]\}$ where

$$\dim(\Theta_{\text{codon}}) = \sum_{j=1}^{M} |\mathcal{C}(y_j)| \tag{25}$$

**Theorem 6** (Codon Profile Convergence Properties) *Under Assumptions 1-3, Codon Profile-constrained optimization using parameter space $\Theta_{codon}$ (main paper Equation 1) and probability function $\Pi_{codon}$ (main paper Equation 2) satisfies:*

1. ***Constraint Preservation**: By construction, codon probabilities $\mathbf{P}_{codon} = \Pi_{codon}(\Theta_{codon}) = softmax(\theta_{j,c}/\sigma)_{c\in\mathcal{C}(y_j)}$ operate only within valid codon space, automatically preserving amino acid identity at all positions $j$*
2. ***Enhanced Convergence**: $\lim_{t\to\infty} \|\nabla F_{CPC}(\Theta_{codon}^{(t)})\| = 0$ with potentially improved conditioning*

*Proof* We establish both constraint preservation and enhanced convergence through dimensionality reduction.

*Step 1: Automatic Constraint Satisfaction.* By construction, Codon Profile uses the probability function $\Pi_{\text{codon}}$ (main paper Equation 2):

$$\mathbf{P}_{\text{codon}} = \Pi_{\text{codon}}(\Theta_{\text{codon}}) = \text{softmax}(\theta_{j,c}/\sigma)_{c\in\mathcal{C}(y_j)} \tag{26}$$

The nucleotide probabilities are obtained via the soft mode transformation $T^{\text{soft}} = R_{\text{nuc}}(\mathbf{P}_{\text{codon}})$ (main paper Equation 8), which reconstructs nucleotide probabilities as weighted sums of codon encodings. Since all operations are restricted to valid codons $\mathcal{C}(y_j)$ for the target amino acid $y_j$, amino acid identity is preserved by construction.

*Step 2: Reduced Parameter Space Analysis.* The optimization operates in the reduced space $\mathbb{R}^{\sum_{j=1}^{M} |\mathcal{C}(y_j)|}$ (dimensionality of $\Theta_{\text{codon}}$) instead of the full RNA parameter space $\mathbb{R}^{L\times 4}$ where $L = 3M$. For standard amino acids, $|\mathcal{C}(y_j)| \in \{1, 2, 3, 4, 6\}$, yielding an average dimensionality reduction by a factor of approximately 4, from $4L$ parameters to $\sum_{j=1}^{M} |\mathcal{C}(y_j)| \approx L$ parameters.

*Step 3: Convergence Rate Improvement.* The constraint eliminates infeasible directions from the gradient flow, potentially improving the condition number of the Hessian matrix. Specifically:

$$\kappa(H_{\mathrm{CPC}}) \leq \kappa(H_{\mathrm{full}}) \tag{27}$$

where $\kappa(\cdot)$ denotes the condition number. This improvement follows from the fact that constraining to a lower-dimensional manifold removes directions of high curvature that are orthogonal to the constraint manifold.

Therefore, convergence is enhanced while automatically maintaining constraint satisfaction. □

##### S5.2.3. Amino Matching Convergence

Amino Matching uses projection-based constraints that maintain differentiability while enforcing amino acid preservation through similarity-based codon selection.

**Codon Probability Function** (main paper Equation 3):

$$\mathbf{P}_{\mathrm{codon}} = \Pi_{\mathrm{amino}}(\Theta^{(j)})_c = \mathrm{softmax}(\langle \Theta^{(j)}, \mathcal{E}[c] \rangle / \sigma)_{c \in \mathcal{C}(y_j)} \tag{28}$$

where $\Theta^{(j)} \in \mathbb{R}^{3\times 4}$ is the RNA parameter matrix for the codon at amino acid position $j$, and $\mathcal{E}[c]$ is the one-hot encoding of codon $c$. The inner product $\langle \Theta^{(j)}, \mathcal{E}[c] \rangle$ measures similarity between RNA parameters and valid codons. The nucleotide probabilities are obtained via $T^{\mathrm{soft}} = R_{\mathrm{nuc}}(\mathbf{P}_{\mathrm{codon}})$ or $T^{\mathrm{hard}} = \mathrm{Onehot}(\mathbf{P}_{\mathrm{codon}})$ (main paper Table 2).

**Theorem 7** (Amino Matching Convergence with Projection) *Under Assumptions 1-3, Amino Matching optimization using projection function $\Pi_{amino}$ (main paper Equation 3) satisfies:*

1. ***Constraint Preservation**: Amino acid constraints are automatically satisfied by restricting codon probabilities to valid codons $\mathcal{C}(y_j)$*
2. ***Full Parameter Space**: Optimization occurs in full RNA parameter space $\mathbb{R}^{L\times 4}$ (where $L = 3M$) while maintaining feasibility*
3. ***Differentiable Projection**: The projection operation $\Pi_{amino}$ preserves gradient flow through inner product similarity*

*Proof* We establish the three key properties through analysis of the differentiable projection mechanism.

*Step 1: Automatic Constraint Satisfaction.* By construction, Amino Matching ensures that codon probabilities are generated only from valid codons through the projection function $\Pi_{\mathrm{amino}}$ (main paper Equation 3):

$$\mathbf{P}_{\mathrm{codon}} = \Pi_{\mathrm{amino}}(\Theta^{(j)})_c = \mathrm{softmax}(\langle \Theta^{(j)}, \mathcal{E}[c] \rangle / \sigma)_{c \in \mathcal{C}(y_j)} \tag{29}$$

Since the softmax is computed only over valid codons $\mathcal{C}(y_j)$ for target amino acid $y_j$, amino acid constraints are automatically preserved. The transformation to nucleotide space via $T^{\mathrm{soft}}$ or $T^{\mathrm{hard}}$ (main paper Table 2) maintains this constraint preservation.

*Step 2: Preservation of Gradient Flow.* The projection function $\Pi_{\mathrm{amino}}$ maintains continuous and well-defined gradients. For the codon probability gradient:

$$\frac{\partial \Pi_{\mathrm{amino}}(\Theta^{(j)})_c}{\partial \Theta^{(j)}} = \frac{\partial}{\partial \Theta^{(j)}} \left[ \mathrm{softmax}(\langle \Theta^{(j)}, \mathcal{E}[c] \rangle / \sigma)_{c \in \mathcal{C}(y_j)} \right] \tag{30}$$

The gradient exists everywhere and is Lipschitz continuous with respect to $\Theta^{(j)}$, ensuring stable optimization dynamics. The full transformation $T^{\mathrm{soft}}$ or $T^{\mathrm{hard}}$ inherits this differentiability through the chain rule.

*Step 3: Full Parameter Space Flexibility.* Unlike Codon Profile, Amino Matching explores the complete RNA parameter space $\mathbb{R}^{L\times 4}$ (where $L = 3M$) while maintaining feasibility through the projection $\Pi_{\mathrm{amino}}$. This provides optimization flexibility while preserving biological constraints, combining the benefits of unconstrained parameter space with automatic constraint satisfaction through geometric projection. □

##### S5.2.4. Lagrangian Method Convergence

The Lagrangian approach [Boyd and Vandenberghe, 2004] uses penalty methods to enforce amino acid constraints through dual variables, with probabilistic codon selection for discrete sequence generation.

**Lagrangian Formulation** (main paper Equation 5):

$$\mathcal{L}_{\mathrm{total}} = f_{\mathrm{model}}(T(\Theta)) + \lambda \cdot \mathcal{C}(\mathbf{P}) \tag{31}$$

where $\lambda \geq 0$ is the Lagrange multiplier that controls penalty strength, and the constraint penalty (main paper Equation 4) measures amino acid constraint violation:

$$\mathcal{C}(\mathbf{P}) = \frac{1}{M} \sum_{j=1}^{M} \min_{c \in \mathcal{C}(y_j)} \|\mathbf{P}^{(j)} - \mathcal{E}[c]\|^2 \tag{32}$$

Here, $\mathbf{P}^{(j)}$ denotes the $3 \times 4$ probability matrix for the codon at amino acid position $j$, $\mathcal{C}(y_j)$ is the set of valid codons encoding amino acid $y_j$, and $\mathcal{E}[c]$ is the one-hot encoding of codon $c$.

**Table 20.** Unified Convergence Summary for ID3 Variants

| Variant | Type | Convergence Target | Key Property |
|---|---|---|---|
| **Det.Soft** | Deterministic Soft | Exact critical point | Strongest guarantee |
| **Sto.Soft** | Stochastic Soft | Expected convergence | Exploration capability |
| **Det.Hard** | Deterministic Hard | $O(B_{\text{STE}})$-neighborhood | Discrete tracking |
| **Sto.Hard** | Stochastic Hard | $O(\sqrt{B_{\text{STE}}^2 + \sigma_G^2})$-neighborhood | Combined benefits |

The discrete sequence selection uses a probabilistic criterion:

$$\text{selected codon} = \arg\max_{c \in \mathcal{C}(y_j)} P(c|\mathbf{P}^{(j)}) \tag{33}$$

where $P(c|\mathbf{P}^{(j)})$ represents the probability of codon $c$ given the probability distribution at position $j$.

**Theorem 8** (Lagrangian Convergence with Standard Subgradient Method) *Under standard regularity conditions, the Lagrangian method with standard subgradient updates [Rockafellar, 1970] for $\lambda$ and probabilistic codon selection converges to points satisfying the KKT conditions:*

$$\begin{aligned} \nabla F(\Theta^*) + \lambda^* \nabla \mathcal{C}(\mathbf{P}^*) &= 0 \\ \mathcal{C}(\mathbf{P}^*) &\leq \epsilon \end{aligned} \tag{34}$$

*with convergence rate $O(1/\sqrt{N})$ for the multiplier $\lambda$.*

*Proof* We establish convergence through standard subgradient analysis with probabilistic selection consistency.

*Step 1: Subgradient Method for $\lambda$ Updates.* Following the main paper (Equation 6), the Lagrange multiplier is updated using the standard subgradient method:

$$\begin{aligned} \lambda^{(t+1)} &= \max(0, \lambda^{(t)} + \eta_t \cdot g^{(t)}) \\ \eta_t &= \eta_0/\sqrt{t+1} \end{aligned} \tag{35}$$

where $g^{(t)}$ is the subgradient of the constraint at iteration $t$, $\eta_t$ is the decaying step size (with $\eta_0$ being the initial step size), and the max operation ensures $\lambda \geq 0$.

The step size satisfies the Robbins-Monro conditions:

$$\sum_{t=1}^{\infty} \eta_t = \infty \quad \text{and} \quad \sum_{t=1}^{\infty} \eta_t^2 < \infty \tag{36}$$

*Step 2: Convergence Rate Analysis.* For the convex constraint penalty function $\mathcal{C}(\mathbf{P})$, the standard subgradient method achieves:

$$\min_{1 \leq t \leq N} \left| \mathcal{C}(\mathbf{P}^{(t)}) - \epsilon \right| \leq \frac{D_{\text{feas}}}{\sqrt{N}} \tag{37}$$

where $D_{\text{feas}}$ is a constant depending on the diameter of the feasible region and the subgradient bound. This establishes the $O(1/\sqrt{N})$ convergence rate.

*Step 3: Probabilistic Selection Consistency.* The discrete sequence selection using $\arg\max_{c \in \mathcal{C}(y_j)} P(c|\mathbf{P}^{(j)})$ ensures that:

- The selected codon is always valid (satisfies amino acid constraints)
- The selection aligns with the continuous optimization objective
- As $\mathbf{P}^{(t)}$ converges, the discrete selection stabilizes

*Step 4: KKT Conditions at Convergence.* At convergence, with $\mathcal{C}(\mathbf{P}^*) \leq \epsilon$ and optimal $\lambda^*$, the stationarity condition holds:

$$\nabla F(\Theta^*) + \lambda^* \nabla \mathcal{C}(\mathbf{P}^*) = 0 \tag{38}$$

The combination of subgradient convergence for $\lambda$ and gradient descent for $\Theta$ ensures convergence to a point satisfying the approximate KKT conditions with tolerance $\epsilon$. □

#### S5.3. Convergence Summary and Hierarchy

##### S5.3.1. Convergence Hierarchy

##### S5.3.2. Practical Implications

**Progressive Complexity**: Each extension adds specific capabilities while maintaining convergence, but also introduces new practical considerations.

**Temperature Tuning**: Lower $\sigma$ improves STE precision and approaches training distribution, but increases Lipschitz constants.

**Variant Selection Guidelines**:

- Use **Det.Soft** when model robustness is high and soft inputs are acceptable
- Use **Det.Hard** when discrete outputs are required and gradient bias is acceptable
- Use **Sto.Soft** when exploration is critical and noise robustness is available
- Use **Sto.Hard** when both exploration and discrete tracking are needed

#### S5.4. Conclusion of Convergence Analysis

This comprehensive convergence analysis establishes theoretical foundations for the ID3 framework across all variants and constraint mechanisms. The three constraint mechanisms (Codon Profile Constraint, Amino Matching Softmax, Lagrangian Multiplier) operate within the unified three-layer architecture while maintaining their distinct approaches to amino acid sequence preservation. The analysis reveals that:

1. All ID3 variants possess well-defined convergence properties under practical assumptions
2. Constraint mechanisms maintain convergence while providing biological sequence preservation
3. The choice of variant should be based on the trade-off between convergence quality and practical requirements
4. Pre-trained model quality fundamentally determines optimization success

These theoretical guarantees provide confidence for practical RNA sequence optimization applications while highlighting the importance of appropriate variant selection based on specific design requirements.

#### S6. CAI Reference Gene Set Construction

##### S6.1. Introduction and Rationale

The Codon Adaptation Index (CAI) serves as a fundamental metric for evaluating the translational efficiency of heterologous genes in bacterial expression systems. Traditional CAI calculations rely on genome-wide codon usage frequencies, which may not accurately reflect the codon preferences of highly expressed genes in specific expression systems. This section details the development and implementation of an expression-system-specific CAI reference set for *E. coli* BL21(DE3), the predominant strain used in recombinant protein production.

##### S6.2. Reference Gene Set Composition

The BL21(DE3) reference gene set was constructed using multiple data sources to ensure comprehensive coverage of highly expressed genes. The final reference set comprises **83 genes** across four functional categories, selected through automated GenBank extraction and validated for biological relevance to protein production.

###### S6.2.1. Ribosomal Proteins (54 genes)

All components of the 30S and 50S ribosomal subunits were included due to their direct role in protein synthesis and consistently high expression levels:

- **30S ribosomal proteins**: rpsA, rpsB, rpsC, rpsD, rpsE, rpsF, rpsG, rpsH, rpsI, rpsJ, rpsK, rpsL, rpsM, rpsN, rpsO, rpsP, rpsQ, rpsR, rpsS, rpsT, rpsU (21 genes)
- **50S ribosomal proteins**: rplA, rplB, rplC, rplD, rplE, rplF, rplI, rplJ, rplK, rplL, rplM, rplN, rplO, rplP, rplQ, rplR, rplS, rplT, rplU, rplV, rplW, rplX, rplY, rpmA, rpmB, rpmC, rpmD, rpmE, rpmF, rpmG, rpmH, rpmI, rpmJ (33 genes)

###### S6.2.2. Translation Factors (11 genes)

Essential components of the translation machinery were selected based on their critical roles in initiation, elongation, and termination:

- **Elongation factors**: fusA (EF-G), tufA (EF-Tu), tufB (EF-Tu), tsf (EF-Ts)
- **Initiation factors**: infA (IF-1), infB (IF-2), infC (IF-3)
- **Release factors**: prfA (RF-1), prfB (RF-2), prfC (RF-3), frr (ribosome recycling factor)

###### S6.2.3. Central Metabolism Enzymes (14 genes)

Key enzymes in glycolysis and associated pathways were included due to their high expression during exponential growth:

- **Primary glycolytic pathway**: pgi, pfkA, fbaA, tpiA, gapA, pgk, gpmA, eno, pykA
- **Associated enzymes**: aceE, aceF, lpdA (pyruvate dehydrogenase complex)
- **Alternative pathway components**: pykF, pfkB

###### S6.2.4. Additional Functional Genes (4 genes)

BL21(DE3)-specific genes important for metabolism and RNA processing:

- fbp (fructose-1,6-bisphosphatase)
- ppsA (phosphoenolpyruvate synthase)
- trmD (tRNA methyltransferase)
- truA (tRNA pseudouridine synthase)

##### S6.3. RSCU-Based CAI Calculation Methodology

###### S6.3.1. RSCU Calculation

For each amino acid family, Relative Synonymous Codon Usage (RSCU) [Sharp and Li, 1987] values are computed from reference sequences:

$$\mathrm{RSCU}_{ij} = X_{ij} \times \frac{n_i}{\sum_k X_{ik}} \tag{39}$$

where $X_{ij}$ is the count of codon $j$ for amino acid $i$ in reference sequences, $n_i$ is the number of synonymous codons for amino acid $i$, and $\sum_k X_{ik}$ is the total count of all codons for amino acid $i$.

###### S6.3.2. Relative Adaptiveness Weights

CAI weights are derived from RSCU values. For each codon $c$ encoding amino acid $i$:

$$w_c = \frac{\mathrm{RSCU}_{ic}}{\max_{c' \in \mathcal{C}(i)}(\mathrm{RSCU}_{ic'})} \tag{40}$$

where $\max_{c' \in \mathcal{C}(i)}(\mathrm{RSCU}_{ic'})$ is the maximum RSCU value among all codons encoding amino acid $i$.

###### S6.3.3. CAI Definition

The Codon Adaptation Index (CAI) for a given sequence $S = (c_1, c_2, \dots, c_M)$ is defined as the geometric mean of the relative adaptiveness values:

$$\mathrm{CAI}(S) = \left(\prod_{j=1}^{M} w_{c_j}\right)^{1/M} \tag{41}$$

where $w_{c_j}$ is the relative adaptiveness weight for codon $c_j$ at position $j$, and $M$ is the total number of codons in the sequence. For numerical stability, this is commonly computed in log-space:

$$\mathrm{CAI}(S) = \exp\left(\frac{1}{M}\sum_{j=1}^{M} \ln(w_{c_j})\right) \tag{42}$$

CAI values range from 0 to 1, with higher values indicating better adaptation to the host organism's codon usage preferences.

###### S6.3.4. Non-Synonymous Codon Exclusion

Critical for standard library consistency, the system excludes non-synonymous codons from CAI calculations:

- **Stop codons**: TAA, TAG, TGA (weight = 0.0)
- **Non-synonymous codons**: ATG (methionine), TGG (tryptophan) are excluded from CAI calculation

##### S6.4. Validation and Comparison

###### S6.4.1. Standard Library Compatibility

The implementation achieves perfect consistency with the Python CAI library through:

- Identical RSCU calculation using the same reference sequence methodology
- Consistent codon exclusion matching non-synonymous and stop codon handling
- Numerical precision maintaining floating-point consistency in geometric mean calculations
- 0% calculation difference with Python CAI library across 96 test sequences

##### S6.5. Complete Codon Weight Tables

The complete 64-codon weight matrices for the BL21(DE3) reference system are available in the supplementary data files. A representative subset is shown below:

**Table 21.** Representative codon weights for BL21(DE3) reference set

| Amino Acid | Codon | RSCU | Weight ($w_i$) |
|---|---|---|---|
| Lysine | AAA | 1.52 | 1.000 |
| Lysine | AAG | 0.48 | 0.316 |
| Leucine | CTG | 3.12 | 1.000 |
| Leucine | CTT | 0.65 | 0.208 |
| Arginine | CGT | 2.45 | 1.000 |
| Arginine | CGC | 1.89 | 0.771 |
| Arginine | CGA | 0.23 | 0.094 |

The BL21(DE3) reference set provides strain-specific codon optimization targets that better reflect the translation machinery preferences in this widely-used expression system, enabling more accurate CAI calculations for recombinant protein production applications.

#### S7. Binary Search for CAI-Aware Discretization

##### S7.1. Problem Setup and Motivation

###### S7.1.1. The CAI-Constrained Optimization Problem

The CAI-aware discretization function (main paper Equation 17) seeks to find the most probable codon sequence under the accessibility-optimized distribution while maintaining sufficient codon adaptation:

$$\begin{aligned}\Psi_{\mathrm{CAI}}^{\mathrm{hard}}(\mathbf{P}_{\mathrm{codon}}) &= \arg\max_{S} P(S|\mathbf{P}_{\mathrm{codon}}) \\ \text{s.t.} \quad \mathrm{CAI}(S) &\geq \text{threshold}\end{aligned} \tag{43}$$

where $S$ is the discrete codon sequence, $\mathbf{P}_{\text{codon}}$ is the accessibility-optimized codon probability distribution from the model, and $P(S|\mathbf{P}_{\text{codon}}) = \prod_{j=1}^{M} P_{\text{codon}_{j,S_j}}$ represents the likelihood of sequence $S$ under this distribution. The CAI threshold threshold is set based on organism-specific requirements (e.g., 0.8 for *E. coli*).

###### S7.1.2. Computational Challenge

This optimization problem is computationally intractable in its general form:

- **Search Space**: The full combinatorial codon space contains $\prod_{j=1}^{M} |\mathcal{C}(y_j)|$ possible sequences, where $\mathcal{C}(y_j)$ is the codon set for amino acid $y_j$ at position $j$
- **Constraint Complexity**: The CAI constraint is non-linear and couples decisions across all positions
- **Greedy Limitations**: Position-wise greedy selection (choosing $\arg\max_c P_{\text{codon}_{j,c}}$ at each position) often violates the CAI constraint

To address this computational challenge, we employ a subspace reduction strategy that restricts the search to a one-dimensional parametric curve. Specifically, we linearly interpolate between two extreme strategies: (1) maximizing sequence probability under the accessibility-optimized distribution $\mathbf{P}_{\text{codon}}$, and (2) maximizing CAI using organism-specific codon frequencies. This dimensional reduction makes the optimization problem tractable while preserving the essential trade-off between accessibility optimization and codon adaptation, as formalized in the following subsection.

##### S7.2. Linear Interpolation Strategy

###### S7.2.1. Interpolation Formulation

We define a parametric family of probability distributions that interpolates between the accessibility-optimized distribution $\mathbf{P}_{\text{codon}}$ and the organism-specific codon usage frequencies $\mathbf{w}$ (main paper Equation 18):

$$\mathbf{P}(\gamma) = \gamma \cdot \mathbf{w} + (1-\gamma) \cdot \mathbf{P}_{\text{codon}} \tag{44}$$

where:

- $\mathbf{w} = [w_{j,c}]$ is the normalized codon usage frequency vector for the target organism
- $\mathbf{P}_{\text{codon}} = [P_{\text{codon}_{j,c}}]$ is the accessibility-optimized distribution from the model
- $\gamma \in [0,1]$ is the interpolation parameter controlling the trade-off:
  - $\gamma = 0$: Pure accessibility optimization (may have low CAI)
  - $\gamma = 1$: Pure CAI optimization (CAI = 1, but may have low model probability)

The reduced optimization problem becomes finding the minimal $\gamma^*$ that satisfies the CAI constraint:

$$\gamma^* = \min\{\gamma \in [0,1] : \text{CAI}(\arg\max \mathbf{P}(\gamma)) \geq \text{threshold}\} \tag{45}$$

where $\arg\max \mathbf{P}(\gamma)$ denotes the sequence obtained by selecting the highest-probability codon at each position under distribution $\mathbf{P}(\gamma)$.

###### S7.2.2. Theoretical Guarantees

The linear interpolation strategy provides strong theoretical guarantees within the one-dimensional search space.

**Theorem 9** (Linear Interpolation Properties) *The linear interpolation strategy satisfies the following properties within the one-dimensional search space* $\mathbf{P}(\gamma) = \gamma \cdot \mathbf{w} + (1-\gamma) \cdot \mathbf{P}_{codon}$ *for* $\gamma \in [0,1]$*:*

1. ***Piecewise Monotonicity:*** *CAI*$(\arg\max \mathbf{P}(\gamma))$ *is piecewise monotonically non-decreasing in* $\gamma$
2. ***Subspace Global Optimality:*** *The solution* $\gamma^*$ *is globally optimal within the one-dimensional interpolation subspace*
3. ***Logarithmic Convergence:*** *Binary search achieves* $O(\log M)$ *convergence on at most* $M$ *switching events*

*Proof* We establish each property through rigorous mathematical analysis.

*Property 1: Piecewise Monotonicity.*

For each amino acid position $j$, the selected codon is determined by:

$$c_j^*(\gamma) = \arg\max_{c \in \mathcal{C}(y_j)} P_{j,c}(\gamma) = \arg\max_{c \in \mathcal{C}(y_j)} [\gamma w_{j,c} + (1-\gamma) P_{\text{codon}_{j,c}}] \tag{46}$$

Consider two codons $c_{\text{acc}}, c_{\text{CAI}} \in \mathcal{C}(y_j)$ where:

- $c_{\text{acc}}$ is the accessibility-optimal codon: $P_{\text{codon}_{j,c_{\text{acc}}}} > P_{\text{codon}_{j,c_{\text{CAI}}}}$
- $c_{\text{CAI}}$ is the CAI-optimal codon: $w_{j,c_{\text{CAI}}} > w_{j,c_{\text{acc}}}$

Define the score difference:

$$\Delta(\gamma) = [\gamma w_{j,c_{\mathrm{CAI}}} + (1-\gamma)P_{\mathrm{codon}_{j,c_{\mathrm{CAI}}}}] - [\gamma w_{j,c_{\mathrm{acc}}} + (1-\gamma)P_{\mathrm{codon}_{j,c_{\mathrm{acc}}}}] \tag{47}$$

This simplifies to:

$$\Delta(\gamma) = \gamma(w_{j,c_{\mathrm{CAI}}} - w_{j,c_{\mathrm{acc}}}) + (1-\gamma)(P_{\mathrm{codon}_{j,c_{\mathrm{CAI}}}} - P_{\mathrm{codon}_{j,c_{\mathrm{acc}}}}) \tag{48}$$

By our definitions, we have:

- $\Delta(0) = P_{\mathrm{codon}_{j,c_{\mathrm{CAI}}}} - P_{\mathrm{codon}_{j,c_{\mathrm{acc}}}} < 0$ (accessibility-optimal codon wins at $\gamma = 0$)
- $\Delta(1) = w_{j,c_{\mathrm{CAI}}} - w_{j,c_{\mathrm{acc}}} > 0$ (CAI-optimal codon wins at $\gamma = 1$)

By the Intermediate Value Theorem, there exists a unique switching point $\gamma_j^{\mathrm{switch}} \in (0,1)$ where codon selection switches from $c_{\mathrm{acc}}$ to $c_{\mathrm{CAI}}$. The derivative $\frac{d\Delta}{d\gamma} = (w_{j,c_{\mathrm{CAI}}} - w_{j,c_{\mathrm{acc}}}) - (P_{\mathrm{codon}_{j,c_{\mathrm{CAI}}}} - P_{\mathrm{codon}_{j,c_{\mathrm{acc}}}}) > 0$ confirms this is the only switching point.

Solving $\Delta(\gamma_j^{\mathrm{switch}}) = 0$ yields:

$$\gamma_j^{\mathrm{switch}} = \frac{P_{\mathrm{codon}_{j,c_{\mathrm{acc}}}} - P_{\mathrm{codon}_{j,c_{\mathrm{CAI}}}}}{(w_{j,c_{\mathrm{CAI}}} - w_{j,c_{\mathrm{acc}}}) + (P_{\mathrm{codon}_{j,c_{\mathrm{acc}}}} - P_{\mathrm{codon}_{j,c_{\mathrm{CAI}}}})} \tag{49}$$

The CAI function for the selected sequence is:

$$\mathrm{CAI}(\gamma) = \exp\left(\frac{1}{M}\sum_{j=1}^{M} \ln(w_{c_j^*(\gamma)})\right) \tag{50}$$

Between switching points, codon selection remains constant, making CAI constant. At each switching point $\gamma_j^{\mathrm{switch}}$, position $j$ switches from the accessibility-optimal codon to the CAI-optimal codon (which has higher weight by definition), causing:

$$\lim_{\gamma \to (\gamma_j^{\mathrm{switch}})^+} \mathrm{CAI}(\gamma) \geq \lim_{\gamma \to (\gamma_j^{\mathrm{switch}})^-} \mathrm{CAI}(\gamma) \tag{51}$$

Therefore, $\mathrm{CAI}(\gamma)$ is piecewise monotonically non-decreasing on $[0,1]$.

*Property 2: Subspace Global Optimality.*

The feasible region is:

$$\mathcal{F} = \{\gamma \in [0,1] : \mathrm{CAI}(\mathbf{P}(\gamma)) \geq \text{threshold}\} \tag{52}$$

By piecewise monotonicity (Property 1), if $\gamma \in \mathcal{F}$, then for any $\gamma' \geq \gamma$, we have $\mathrm{CAI}(\gamma') \geq \mathrm{CAI}(\gamma) \geq$ threshold, implying $\gamma' \in \mathcal{F}$. Therefore, $\mathcal{F} = [\gamma^*, 1]$ where $\gamma^* = \inf \mathcal{F}$ (assuming feasibility).

The optimization problem within the interpolation subspace is:

$$\min_{\gamma \in [0,1]} \gamma \quad \text{subject to} \quad \mathrm{CAI}(\mathbf{P}(\gamma)) \geq \text{threshold} \tag{53}$$

Since the objective $f(\gamma) = \gamma$ is linear (hence convex) and the feasible region $\mathcal{F} = [\gamma^*, 1]$ is convex, the global optimum within the subspace is $\gamma^*$, which minimizes $\gamma$ (maximizes consistency with accessibility-optimized distribution) while satisfying the CAI constraint.

*Property 3: Logarithmic Convergence.*

Let $\mathcal{G} = \{0 = \gamma_0 < \gamma_1 < \cdots < \gamma_K = 1\}$ be the set of all switching points (at most $M$ switching events, one per position). The binary search operates on this discrete set.

At each iteration, the search interval is halved. To locate $\gamma^*$ within tolerance $\varepsilon$, we need:

$$\frac{1}{2^k} \leq \varepsilon \Rightarrow k \geq \log_2(1/\varepsilon) \tag{54}$$

For discrete search on at most $M$ switching events, the complexity is $O(\log M)$ iterations. □

Remark 1 (Search Space Limitations) *The binary search algorithm explores only a one-dimensional subspace of the full combinatorial codon space. Specifically:*

- ***Full Space Size:*** $\prod_{j=1}^{M} |\mathcal{C}(y_j)|$ *possible codon combinations*
- ***Explored Space:*** *The 1D parametric curve* $\mathbf{P}(\gamma)$ *for* $\gamma \in [0,1]$
- ***Reachable Codons:*** *At position* $j$*, only codons with* $w_{j,c} > 0$ *OR* $P_{codon_{j,c}} > 0$ *can be selected*
- ***Potential Gap:*** *Codon combinations outside the interpolation path are not explored*

*While this linear interpolation captures the primary accessibility-CAI trade-off axis, alternative codon combinations outside this path may exist. The algorithm provides a computationally efficient approach with strong guarantees within the explored subspace, representing a pragmatic balance between theoretical optimality and practical tractability.*

##### S7.3. Implementation

The binary search algorithm exploits the piecewise monotonicity property (Theorem 9, Property 1) to efficiently find the minimal $\gamma^*$ satisfying the CAI constraint. To enhance diversity and explore sequences beyond the interpolation subspace, we employ a greedy refinement strategy with hash-based duplicate detection. This combination ensures both theoretical optimality within the interpolation subspace and practical sequence diversity.

#### S8. CAI Integration: Penalty vs. Direct Approach

This section compares two implementation approaches for integrating CAI constraints into the ID3 framework: penalty-based optimization (multi-objective joint loss during gradient descent) and direct STE approach (post-optimization binary search). These two approaches are not mutually exclusive and can be combined: the penalty-based approach can be applied during optimization to guide the model toward better CAI values, followed by the direct approach for fine-tuning CAI targets through post-processing. Statistical performance comparison is provided in Section S2.5.3. We present the mathematical formulations and design rationale for selecting the direct STE approach as the default method in the main framework.

##### S8.1. Two Implementation Approaches

**Direct STE Approach (Hard Mode Only).** The direct approach uses the straight-through estimator without penalty terms during optimization, focusing solely on accessibility optimization:

$$\min_{\Theta} \mathcal{L}_{\text{Access}}(T(\Theta)) \tag{55}$$

This is followed by post-optimization binary search enhancement to achieve target CAI values. This method is only applicable to hard discretization mode, as the binary search requires discrete sequences to compute CAI and perform interpolation. The approach separates accessibility optimization from CAI adjustment, with CAI targets achieved through post-processing binary search interpolation between accessibility-optimal and CAI-optimal codon distributions, as detailed in Section S7.

**Penalty-Based Approach (Both Soft and Hard Modes).** The penalty-based approach incorporates CAI constraints directly into the optimization objective through a unified loss function:

$$\min_{\Theta} \mathcal{L}_{\text{Access}}(T(\Theta)) + \lambda_{\text{CAI}}\mathcal{L}_{\text{CAI}}(\Theta) \tag{56}$$

where $\mathcal{L}_{\text{CAI}}(\Theta) = (\text{ECAI}(T(\Theta)) - \text{CAI}_{\text{target}})^2$ is computed using the Expected CAI. This approach works with both soft mode (continuous probability distributions) and hard mode (discrete sequences), making it the only option for CAI optimization in soft discretization mode. For hard mode, this approach can be used alone or combined with the direct STE approach for enhanced CAI control. The penalty coefficient $\lambda_{\text{CAI}}$ is updated using the same subgradient method as the Lagrangian multiplier in the Lagrangian constraint mechanism (main paper Equation 6), enabling adaptive penalty strength adjustment during optimization.

**Expected CAI (ECAI):** For the penalty-based optimization approach, we define the Expected CAI that supports both continuous probability sequences and discrete one-hot sequences:

$$\text{ECAI}(\mathbf{P}_{\text{codon}}) = \left( \prod_{j=1}^{M} \sum_{c \in \mathcal{C}(y_j)} P_{\text{codon}_{j,c}} \cdot w_c \right)^{1/M} \tag{57}$$

where $\mathbf{P}_{\text{codon}} = (P_{\text{codon}_1}, P_{\text{codon}_2}, \ldots, P_{\text{codon}_M})$ represents either continuous probability distributions or discrete one-hot sequences at each position $j$, $\mathcal{C}(y_j)$ is the set of valid codons for amino acid $y_j$, and $w_c$ is the CAI weight for codon $c$. This unified formulation enables ECAI computation from both probabilistic constraint outputs and discrete sequences, supporting gradient-based optimization of the penalty term.

##### S8.2. Performance Equivalence and Design Choice

Complete results for both approaches across 12 variants and 12 proteins are presented in Tables S2–S3 and detailed statistical comparison in Section S2.5.3. The use of penalty-based optimization during training shows minimal performance difference compared to training without penalty (mean difference = 0.005 kcal/mol, p = 0.661, Cohen's d = 0.012). Notably, the best-performing variant (Amino.Sto.Hard +Penalty, rank 1 in Table S2) does employ the penalty-based approach during optimization.

The superior performance of the direct STE approach with hard discretization can be attributed to two key factors. First, hard discretization enables the pre-trained DeepRaccess model to operate on discrete sequences matching its training distribution, avoiding the distribution shift inherent in soft probability inputs. Second, the post-optimization binary search allows precise CAI targeting without the gradient noise and optimization conflicts introduced by multi-objective penalty terms during training. These technical advantages explain why hard mode variants dominate the top rankings in Table S2, with the direct approach achieving the optimal balance between accessibility optimization and CAI constraint satisfaction.

#### S9. UTR Template Specifications

The ID3 framework employs fixed 5' and 3' UTR sequences derived from the pET-11a expression vector (Novagen), optimized for T7 RNA polymerase-based protein expression in *E. coli* [Studier and Moffatt, 1986].

**5' UTR (70 nucleotides):**

```
GGGAATTGTGAGCGGATAACAATTCCCCTCTAGAAATAATTTTGTTTAACTTTAAGAAGGAGATATACAT
```

**3' UTR (63 nucleotides):**

GATCCGGCTGCTAACAAAGCCCGAAAGGAAGCTGAGTTGGCTGCTGCCACCGCTGAGCAATAA

Vector details: https://www.snapgene.com/plasmids/pet_and_duet_vectors_(novagen)/pET-11a

#### References

Y. Bengio, N. Léonard, and A. Courville. Estimating or propagating gradients through stochastic neurons for conditional computation. *arXiv preprint arXiv:1308.3432*, 2013.

S. Boyd and L. Vandenberghe. *Convex Optimization*. Cambridge University Press, Cambridge, UK, 2004. doi: 10.1017/CBO9780511804441.

E. Jang, S. Gu, and B. Poole. Categorical reparameterization with gumbel-softmax. arXiv preprint arXiv:1611.01144, 2016.

Y. Nesterov. *Introductory Lectures on Convex Optimization: A Basic Course*, volume 87 of *Applied Optimization*. Springer, 2004. doi: 10.1007/978-1-4419-8853-9.

R. T. Rockafellar. *Convex Analysis*, volume 28 of *Princeton Mathematical Series*. Princeton University Press, Princeton, NJ, 1970.

P. M. Sharp and W. H. Li. The codon adaptation index-a measure of directional synonymous codon usage bias, and its potential applications. *Nucleic Acids Research*, 15(3):1281–1295, 1987.

F. W. Studier and B. A. Moffatt. Use of bacteriophage t7 rna polymerase to direct selective high-level expression of cloned genes. *Journal of Molecular Biology*, 189(1):113–130, 1986.
